# Engaging in word recognition elicits highly specific modulations in visual cortex

**DOI:** 10.1101/2022.10.10.511419

**Authors:** Alex L. White, Kendrick Kay, Kenny Tang, Jason D. Yeatman

## Abstract

A person’s cognitive state determines how their brain responds to visual stimuli. The most common such effect is a response enhancement when stimuli are task-relevant and attended rather than ignored. In this fMRI study, we report a surprising twist on such attention effects in the visual word form area (VWFA), a region that plays a key role in reading. We presented participants with strings of letters and visually similar shapes which were either relevant for a specific task (lexical decision or gap localization) or ignored (during a fixation dot color task). In the VWFA, the enhancement of responses to attended stimuli occurred only for letter strings, whereas the shapes evoked *smaller* responses when attended than when ignored. The enhancement of VWFA activity was accompanied by strengthened functional connectivity with higher-level language regions. These task-dependent modulations of response magnitude and functional connectivity were specific to the VWFA and absent in the rest of visual cortex. We suggest that language regions send targeted excitatory feedback into the VWFA only when the observer is trying to read. This feedback enables the discrimination of familiar and nonsense words, and is distinct from generic effects of visual attention.

## INTRODUCTION

Visual cortex is capable of processing a wide variety of stimuli for any number of behavioral tasks. This raises a question: how exactly does the specific information required at any given moment get selected and used to execute a task? One key finding is that visual cortex does not perform a static stimulus-response mapping. Rather, the organism’s goals influence ongoing activity ^1,2^. The most prominent forms of top-down influence relate to attention: stimuli that are relevant to the current task evoke stronger responses than stimuli that are irrelevant, due to selection on the basis of visual field location or non-spatial features ^3–5^. However, we hypothesize that the brain performs more than simple amplification of relevant stimuli, because different tasks require particular visual information to be routed to different brain networks.

The focus of this study is word recognition, an important visual task that engages a specific network beyond visual cortex ^6,7^. The “visual word form area” (VWFA) in left ventral occipito-temporal cortex is key to the transformation of retinal input into lexical information that is conveyed to language regions during reading ^8–11^. Neighboring regions are specialized for recognizing faces, bodies, objects and scenes ^12^.

There are two ways of conceptualizing the VWFA, which are not mutually exclusive: first, it could essentially be a visual area, with intrinsic selectivity for certain stimulus features in certain parts of visual space. Second, the VWFA could be unique due to its connection to the brain’s language network, which regulates its activity and infuses it with lexical information via top-down signals.

Some evidence favors the first view. Although the VFWA responds most strongly to strings of letters, it does respond in a graded fashion to other categories of images ^13–15^. Importantly, the VWFA has several subregions that increase in selectivity along the posterior-to-anterior axis ^11,16–19^. It is also sensitive to the visual field position ^20^, responding most strongly to words in the fovea and right parafovea ^21,22^. Lastly, there are spatial attention effects in the VWFA similar to those elsewhere in visual cortex ^3^: words evoke stronger responses when they are at an attended location than an ignored location ^14,18^.

The VWFA’s function extends beyond the purely visual, however. Some authors argue that it represents whole words as distinct identities ^23–25^. It also responds differentially to frequent words, infrequent words, and novel pseudowords ^11,26^. The VWFA’s sensitivity to such higher-level linguistic features can be modulated by the participant’s task ^11,15,27^. Moreover, Braille reading ^28–30^ and certain auditory judgments ^31–34^ have been reported to activate the VWFA without visual stimulation. (Note that some similar cognitive effects also occur in other category-selective visual areas ^35,36^.)

Altogether, these extra-visual and cognitive effects have led some researchers to the second view: that the VWFA plays a special role in reading only as a result of feedback from the language network ^37^. Indeed, the VWFA has white matter connections to spoken language regions, as well as regions associated with attentional control ^38–41^. There is also strong resting state functional connectivity between the VWFA and those regions ^38,42–45^ (see also ^46^). However, the effects of such connectivity on the VWFA’s activity remain unspecified, and connectivity has not been linked to task effects on stimulus-driven responses.

The goal of this study is to clarify the functional properties of the VWFA by measuring its stimulus selectivity and functional connectivity under varying task demands. We asked: what is the precise nature of the interaction between bottom-up stimulus features and top-down modulation? Does “visual attention” boost any task-relevant sensory signal in the VWFA, or are top-down enhancements contingent on engagement in a linguistic task? Do task effects in the VWFA differ from those in other visual areas? To answer these questions, we recorded fMRI activity while participants viewed words and non-letter shapes and performed three different tasks in which those stimuli were either relevant and attended or irrelevant and ignored.

On each trial, one stimulus appeared at a random position along the horizontal meridian (**Figure 1A**). Each stimulus was either a string of four letters (forming either a real word or a pronounceable pseudoword), or a string of four squares and circles matched in size to the letters (polygons are a good stimulus to drive the VWFA^14^).

**Figure 1.**
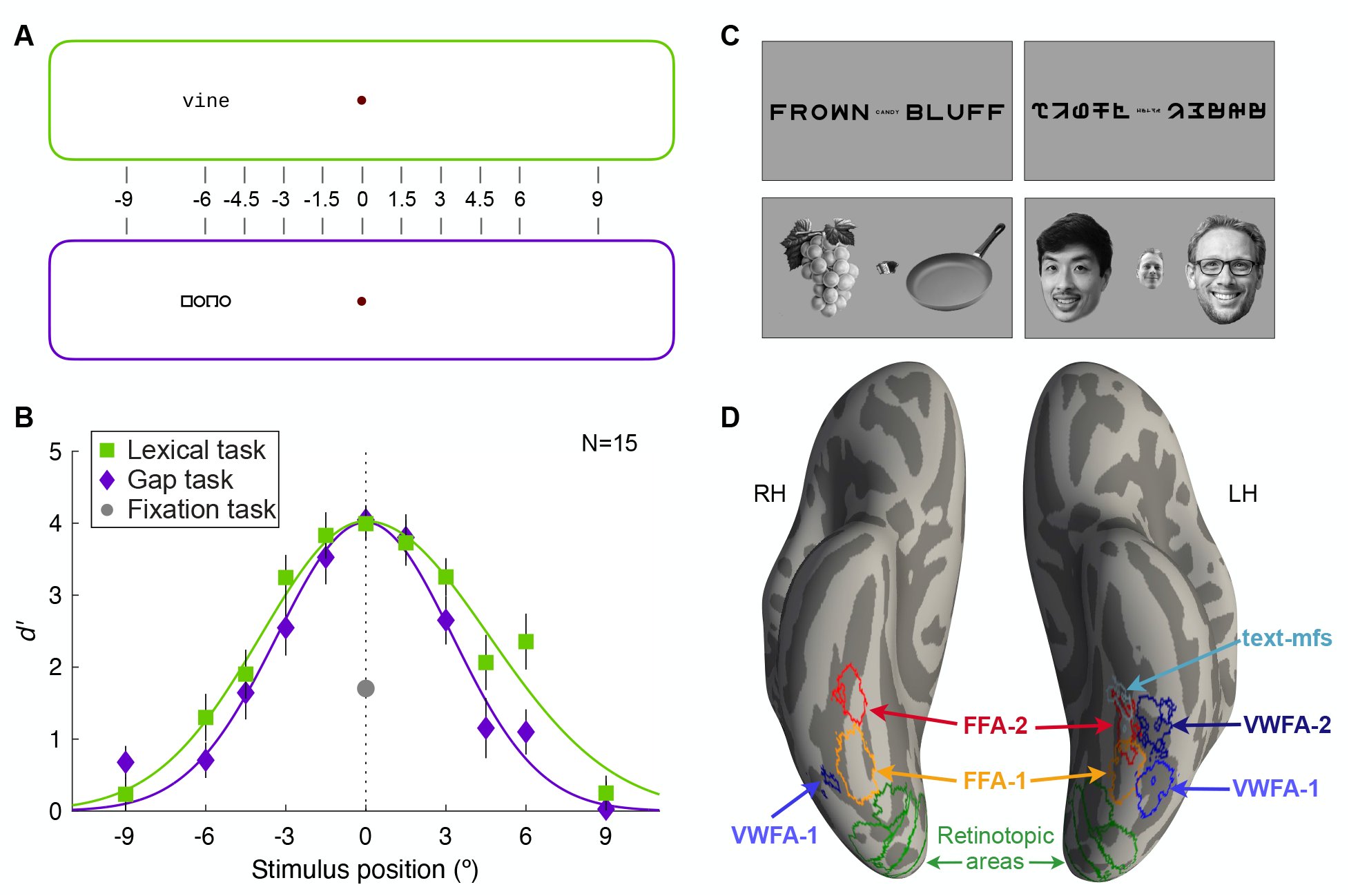
Stimuli, behavioral performance, and ROIs. **(A)** Two example stimuli, with the 11 possible stimulus locations marked in degrees of visual angle. **(B)** Mean behavioral accuracy (*d’*). See also Figure S1. **(C)** Examples of the stimulus categories in the localizer scan. Faces of people other than the three authors, shown here, were actually used. **(D)** Most likely ROI locations on a ventral view of an average cortical surface (Freesurfer’s “fsaverage”). We first projected individual ROIs onto the fsaverage surface. For each ROI, the outline shown here is the half-max contour of the count of participants who had that ROI at each vertex.

In half the scans, participants attended to the stimuli while maintaining gaze fixation on a central dot. When they saw letter strings, they performed the *lexical decision task*: to report whether the stimulus was a real word or a pseudoword. When they saw shape strings, they performed the *gap localization task*: to report whether a gap in one of the inner two shapes was on the top or bottom side. (In a separate behavioral experiment, we found that performance in these two tasks varies similarly with stimulus eccentricity. Therefore, they are well matched in many visual properties). In the other half of scans, participants ignored the letters and shapes and performed the *fixation color detection task*: to report whether or not the fixation dot turned slightly red. The visibility of the red color was controlled by a staircase to maintain a consistent level of difficulty.

We designed these tasks to strictly control attention. During the fixation task, spatial attention had to be focused tightly on the fixation dot. The dot changed color simultaneously with the appearance of the letters or shapes, which lasted for only 150 ms. The participant did not have time to switch attention or make a saccade from the fixation dot to the stimuli. During both the gap task and the lexical task, the participant was motivated to widen their spatial attention to perceive a stimulus at any location. Although the participant had little time to focus attention onto any one stimulus before it disappeared, they may have been able to prioritize its sensory memory trace, thereby boosting the subsequent BOLD response. Regardless, the spatial attention manipulation is matched for the letters and shape stimuli.

In sum, differences in the BOLD response to letter and shape strings during the fixation task can be primarily attributed to neuronal selectivity for stimulus features. Differences between the gap and lexical tasks in terms of how the BOLD response is elevated or suppressed relative to their fixation-task baselines can be attributed to what information the participant is instructed to judge, and how that judgement matches the function of each brain region. A generic attentional enhancement, as is common in visual cortex (including face- and word-selective regions^14,47^), would predict larger responses during both the lexical and gap tasks compared to the fixation task.

## RESULTS

### Task performance

In both the gap task and the lexical task, discrimination accuracy (*d’;* **Figure 1B)** was near perfect for stimuli at fixation (0º), but dropped off quickly with increasing eccentricity, approaching chance (<55% correct) by ±9º eccentricity. Averaged across stimulus positions, the lexical, gap, and fixation tasks were similarly difficult: mean (± SEM) *d’* = 1.76 ± 0.15, 1.35 ± 0.11, and 1.71 ± 0.15, respectively. *d’* in the gap task was slightly lower than in the lexical task (t(14)=2.14, *P*=0.05, BF=1.53), and than in the fixation task (t(14)=2.82, *P*=0.014; BF=4.34). The lexical and fixation tasks did not differ (t(14)=0.22, *P*=0.83; BF=0.27). We fit a linear mixed-effect (LME) model to predict *d’* as a function of task (lexical vs. gap), absolute eccentricity, and hemifield (left vs. right). Only the effect of eccentricity was significant (F(1,313)=49.7, *P*=10^−10^), and it did not interact with task (F(1,313)=1.74, *P*=0.19). For performance separately for real and pseudowords, see **Figure S1**.

### A surprising interaction of stimulus and task effects in the left VWFAs

We used independent localizer scans (**Figure 1C**) to define regions of interest (ROIs) on each participant’s cortical surfaces (**Figure 1D**). We focus on two text-selective regions in left occipito-temporal sulcus, VWFA-1 and VWFA-2 ^18^. Also of interest are two face-selective regions, FFA-1 and FFA-2, medial to the text-selective regions in both hemispheres, as well as posterior retinotopic visual areas (V1-hV4, VO1/2 and LO1/2).

The mean responses in left VWFA-1 and VWFA-2 are shown in **Figure 2**: beta weights from a GLM in units of percent signal change (p.s.c.) relative to fixation on a blank screen. Both areas had larger responses to letter strings than shape strings (both F>23, p<10^−5^). They also preferred stimuli in the central visual field ^21^: responses decreased with absolute eccentricity (both F>16, *P*<10^−4^). The overall preference for letters did not vary with eccentricity (VWFA-1: *P*=0.42, BF=0.012; VWFA-2: *P*=0.99, BF=0.009). VWFA-1 responded more strongly to stimuli in the right visual field than the left (F(1,584)=13.9, *P*=2×10^−4^, BF=782). That hemifield asymmetry was numerically larger for letters than shapes but did not significantly interact with stimulus type (BF=0.6) or task (BF=0.21). VWFA-2 responded equally to stimuli in left and right hemifields (F<1, BF=0.14).

**Figure 2:**
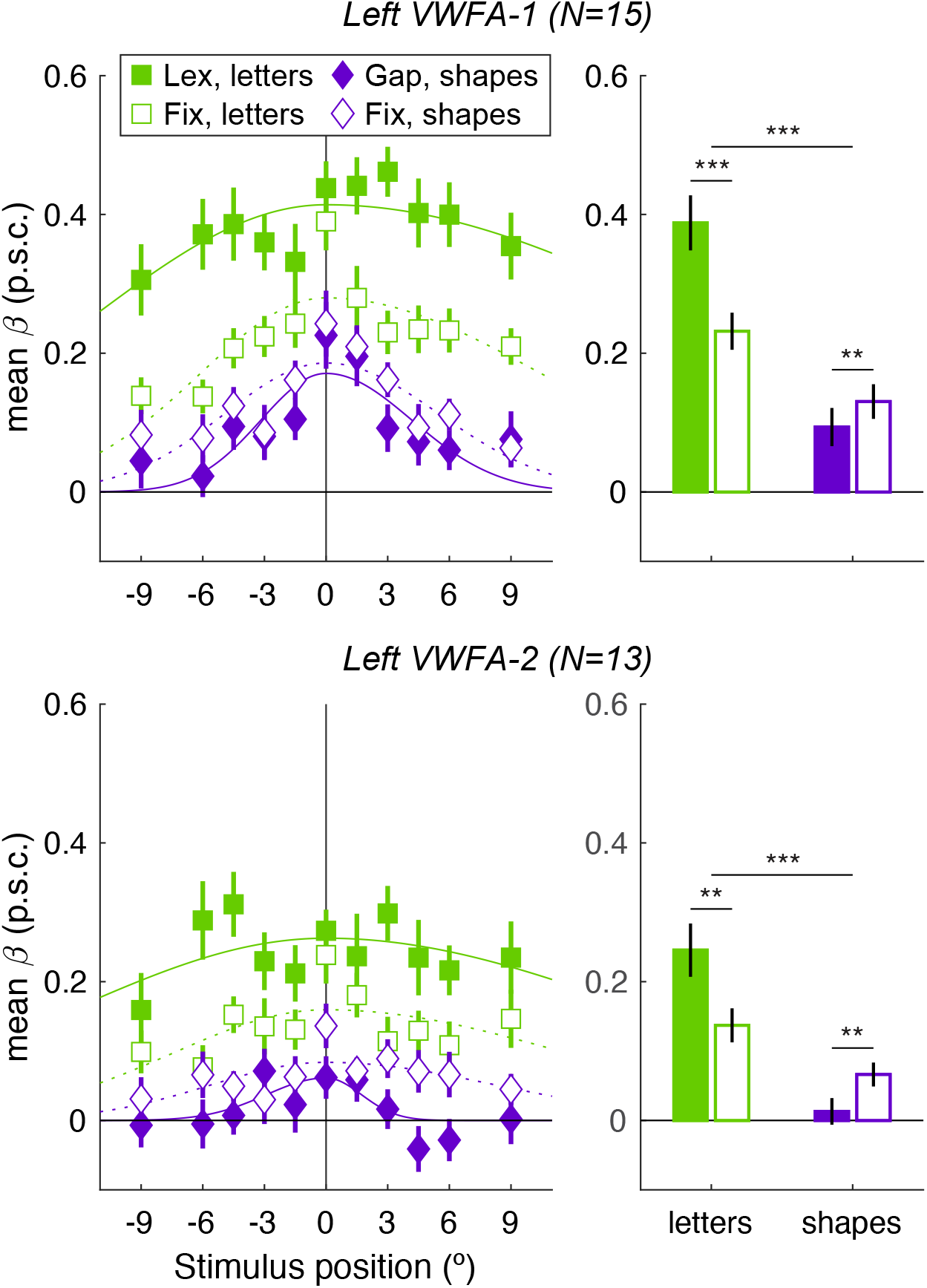
Mean BOLD responses in two sub-regions of the VWFA. *Left column:* Responses as a function of stimulus position, stimulus type, and task (Lex = lexical task; Fix = fixation task). The smooth lines are asymmetric Gaussian functions fit to each condition. *Right column:* The same responses collapsed across stimulus positions. Asterisks indicate the *P*-value for the task effects (short bars over each stimulus type), and task-by-stimulus interactions (long bars): *** *P*<0.001; ** *P*<0.01.

We also found that letter strings evoked much larger VWFA responses during the lexical decision task (when task-relevant) than during the fixation task (when ignored; both sub-regions’ *P*<0.01; BF>14). The ratio of mean lexical task response to mean fixation task response was 1.67 in VWFA-1 and 1.77 in VWFA-2. The task effect did not vary with stimulus eccentricity (VWFA-1: *P*=0.13, BF=0.03; VWFA-2: *P*=0.49, BF=0.01).

However, the non-letter shape strings evoked *smaller* responses when they were task-relevant (during the gap task) than when they were ignored (during the fixation task). The mean ratio of gap task response to fixation task response was 0.72 in VWFA-1 and 0.18 in VWFA-2. This task-related *suppression* was statistically reliable in both areas (*Ps*<0.01, BFs>15). Importantly, there were also significant interactions between stimulus type (letters, shapes) and task (“attend-stimuli”, “attend-fixation”). In VWFA-1, t(14)=9.51, *P*=2×10^−7^; BF=8×10^4^. In VWFA-2, t(12)=4.05, *P*=0.002; BF=27.3. For an analysis of how activity varied across trials within each block, see **Supplementary Figure S2**.

These curious task effects were largely absent in the other visual ROIs we analyzed. **Figure 3** plots mean responses in other areas, collapsed over stimulus position, and **Table 1** lists the statistics. **Supplementary Figure S2A** visualizes the task-by-stimulus interaction on the average cortical surface, showing that it is restricted to the VWFAs. **Supplementary Figure S3** shows data from a third text-selective area, text-mfs.

**Table 1.** Statistics on task effects and their interaction with stimulus type on BOLD responses. Under “Hem”, B = both hemispheres together, L = left, R = right. SE = standard error of the mean, *P =* t-test p-value, corrected for false discovery rate across the 15 ROIs; Significant values are in bold.

| ROI | Hem | N | Task effect for Letters<br>(Lex. - Fix.) |  |  | Task effect for Shapes<br>(Gap - Fix.) |  |  | Interaction (Lex.-Fix.) -<br>(Gap-Fix.) |  |  |
| --- | --- | --- | --- | --- | --- | --- | --- | --- | --- | --- | --- |
|  |  |  | Mean (SE) | P | BF | Mean (SE) | P | BF | Mean (SE) | P | BF |
| V1 | B | 15 | 0.01 (0.02) | 0.90 | 0.3 | -0.01 (0.02) | 0.79 | 0.3 | 0.02 (0.02) | 0.87 | 0.24 |
| V2 | B | 15 | 0.02 (0.02) | 0.63 | 0.4 | 0.02 (0.02) | 0.33 | 0.5 | -0.00 (0.04) | 0.87 | 0.27 |
| V3 | B | 15 | 0.03 (0.01) | 0.08 | 1.5 | 0.03 (0.02) | 0.15 | 1.4 | -0.00 (0.02) | 0.87 | 0.23 |
| hV4 | B | 15 | 0.04 (0.01) | 0.06 | 3.6 | 0.04 (0.03) | 0.26 | 0.7 | 0.00 (0.03) | 0.87 | 0.24 |
| VO | B | 15 | 0.02 (0.02) | 0.82 | 0.3 | 0.03 (0.01) | 0.09 | 2.0 | -0.02 (0.02) | 0.57 | 0.45 |
| LO | B | 15 | 0.00 (0.01) | 0.82 | 0.3 | **0.03 (0.01)** | **0.05** | **5.8** | -0.02 (0.01) | 0.34 | 0.61 |
| VWFA-1 | L | 15 | **0.16 (0.02)** | **10<sup>-4</sup>** | **10<sup>4</sup>** | **-0.04 (0.01)** | 0.01 | **16** | **0.19 (0.02)** | **10<sup>-6</sup>** | **10<sup>6</sup>** |
| VWFA-1 | R | 11 | 0.08 (0.03) | 0.08 | 2.0 | 0.02 (0.02) | 0.59 | 0.4 | 0.07 (0.05) | 0.34 | 0.78 |
| VWFA-2 | L | 13 | **0.11 (0.03)** | **0.01** | **14** | **-0.05 (0.01)** | 10<sup>-2</sup> | **39** | **0.16 (0.04)** | **10<sup>-2</sup>** | **76** |
| text-mfs | L | 9 | **0.08 (0.02)** | **0.01** | **15** | -0.03 (0.01) | 0.19 | 1.1 | **0.11 (0.03)** | **0.03** | **99** |
| FFA-1 | L | 14 | **0.03 (0.01)** | 10<sup>-2</sup> | **51** | -0.01 (0.01) | 0.33 | 0.6 | **0.05 (0.02)** | **0.04** | **7.7** |
| FFA-1 | R | 15 | 0.00 (0.01) | 0.90 | 0.3 | 0.00 (0.01) | 0.74 | 0.3 | -0.00 (0.02) | 0.87 | 0.23 |
| FFA-2 | L | 13 | 0.01 (0.02) | 0.82 | 0.3 | -0.00 (0.02) | 0.92 | 0.3 | 0.01 (0.03) | 0.87 | 0.25 |
| FFA-2 | R | 12 | 0.00 (0.01) | 0.82 | 0.3 | 0.01 (0.02) | 0.64 | 0.4 | -0.01 (0.03) | 0.87 | 0.33 |
| Broca's | L | 15 | **0.08 (0.01)** | 10<sup>-3</sup> | **380** | **-0.04 (0.01)** | 10<sup>-2</sup> | **26** | **0.12 (0.02)** | 10<sup>-3</sup> | 10<sup>5</sup> |

**Figure 3:**
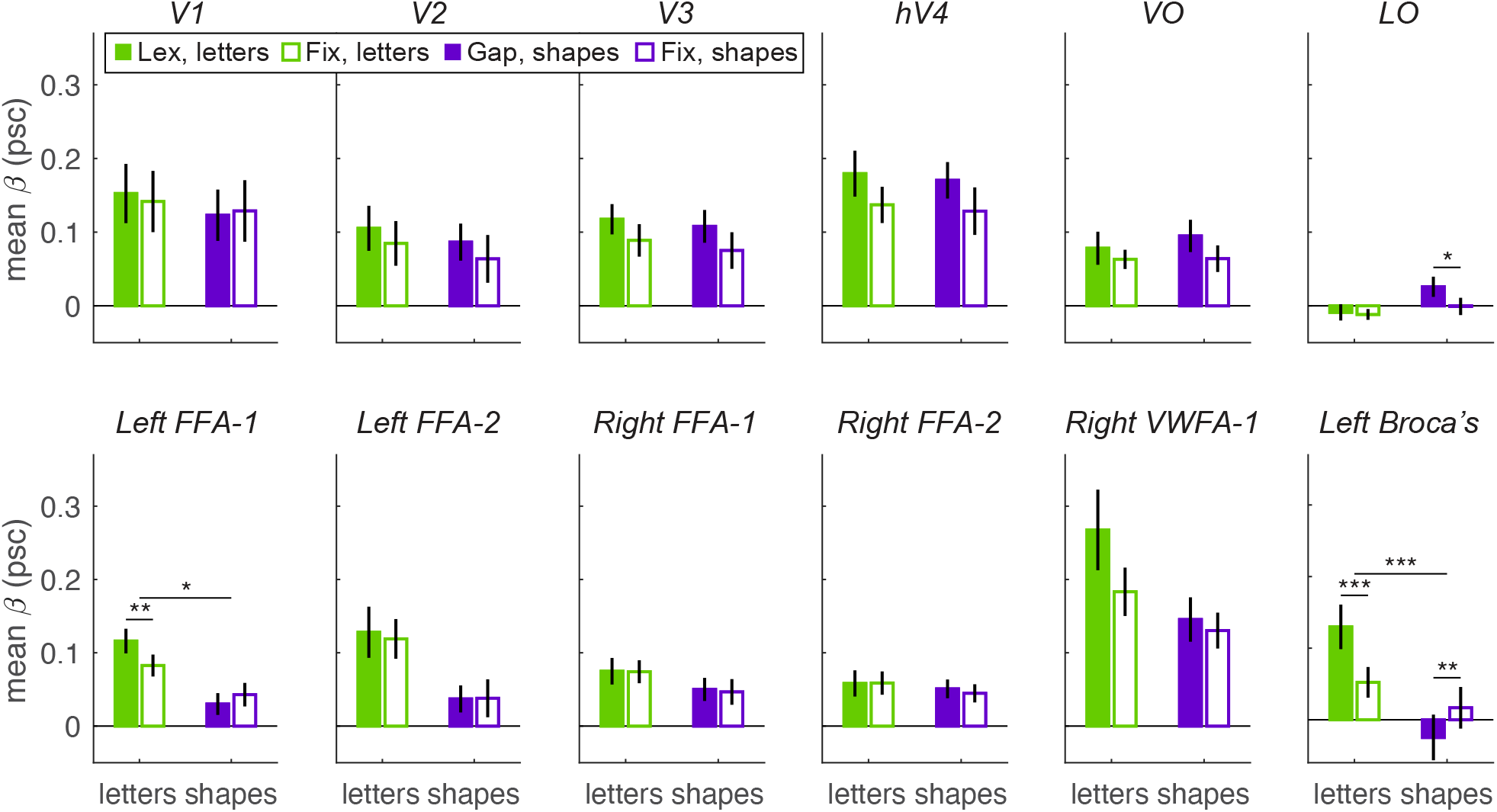
Mean BOLD responses, collapsed across stimulus position, in other areas. The top row is for retinotopic areas, collapsed across left and right hemispheres. Within V1-hV4 we extracted the response on each trial from vertices that had PRFs centered over that stimulus position (excluding ±9º, which was not localizable). Areas VO and LO were defined from published atlases and responses were averaged over all vertices within them. Abbreviations and significance stars as in Figure 2.

Responses in retinotopic visual areas (V1-LO) were weak overall, perhaps because of the brief duration and small size of the stimuli, despite selecting subsets of voxels for each position. Nonetheless, main effects of task-relevance (stimuli attended>ignored) emerged in V3, hV4, and VO (only hV4 survived correction for false discovery rate). This could be an effect of spatial attention generically boosting responses. The only area with an overall preference for non-letter shapes was LO (*P*=0.005, BF=26). LO responded more strongly to shapes during the gap task than during the fixation task (the opposite of what we observed in the VWFAs).

One face-selective area, left FFA-1, also responded more strongly to letters during the lexical than the fixation task, but none of the other three face areas showed any task effects or interactions with stimulus type. All face-selective areas responded more strongly to letters than shapes (all *P*<0.001, BF>20), except right FFA-2 (BF=0.26). Right VWFA-1 responded most strongly to letters during the lexical task, but the task-by-stimulus interaction was not significant (Table 1).

Finally, we also defined a putative left frontal Broca’s area (illustrated in **Figure 5A**) by contrasting (in the average brain) responses to letters vs. shapes, across tasks and stimulus positions. This area was at the border of the left ventral precentral sulcus and inferior frontal sulcus, overlapping Broadman’s Area 44 and the pars opercularis, a region known to play a role in lexical decision ^48,49^. Activity in this Broca’s area showed the same task-by-stimulus interaction as the VWFAs: much larger response to letters in the lexical than fixation task, but smaller (even negative) responses to shapes in the gap than fixation task (bottom right panel of Figure 3). For activity as a function of stimulus position, see Figure 4 and **Supplementary Figure S2D**.

**Figure 4:**
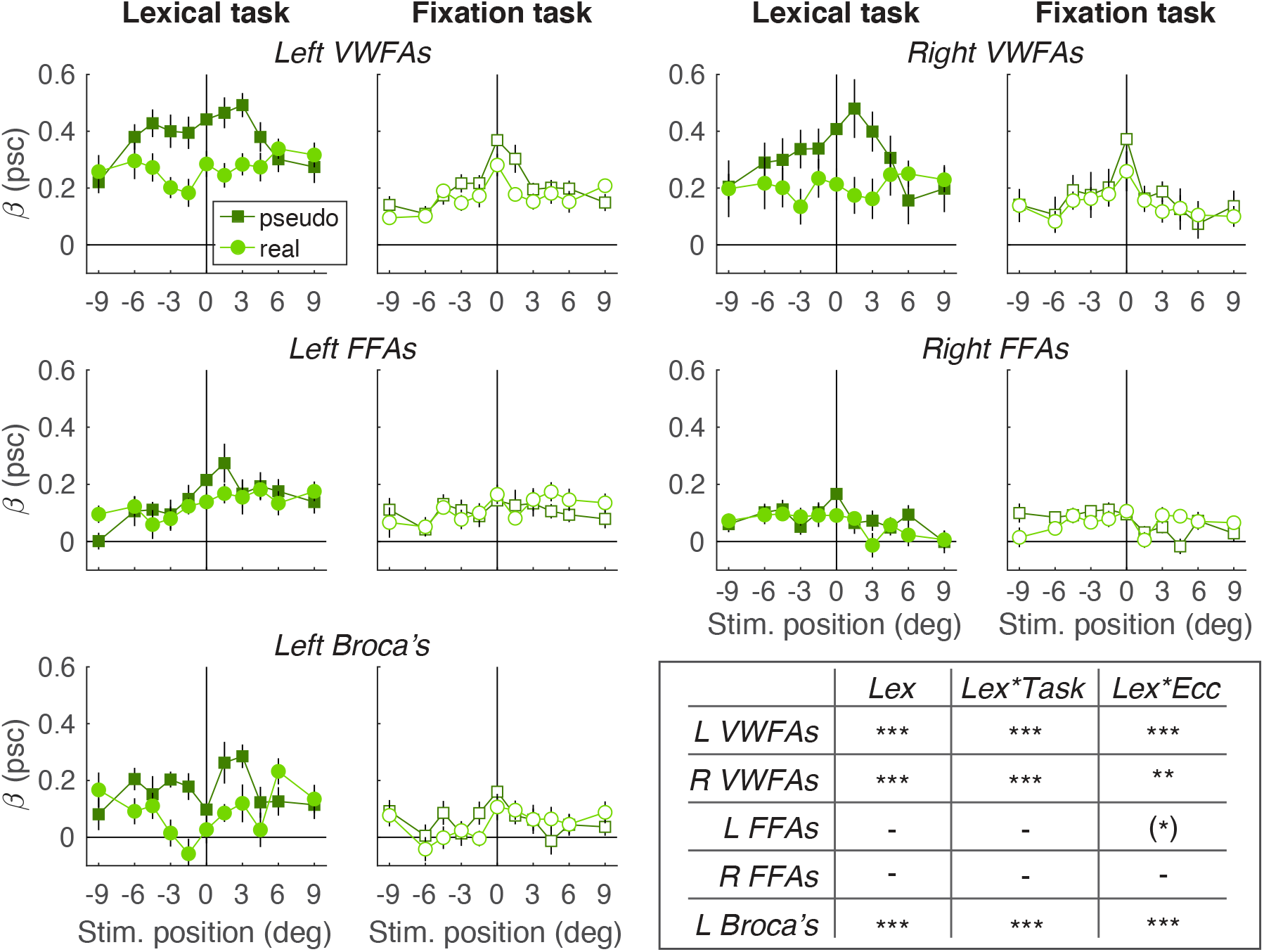
Lexicality effects. Mean BOLD responses to pseudowords (dark green squares) vs. real words (light green circles) in five brain areas. “VWFAs” and “FFAs” are the average over the two subregions of each category-selective area. For each area, the left plot shows data from the lexical decision task, and the right from the fixation task. Error bars = +/- 1 SEM. The table summarizes LME models that predict the response as a function of these factors: “Lex”, the main effect of lexicality; “Lex*Task”, the interaction of lexicality and task, and “Lex*Ecc”, the interaction of lexicality and absolute stimulus eccentricity. *P*-values are corrected for false discovery rate: \*\*\**P*<0.001, \*\**P*<0.01, \**P*<0.05. One asterisk for left FFAs is in parentheses because the Bayes factor favored the null hypothesis (BF=0.16).

**Figure 5:**
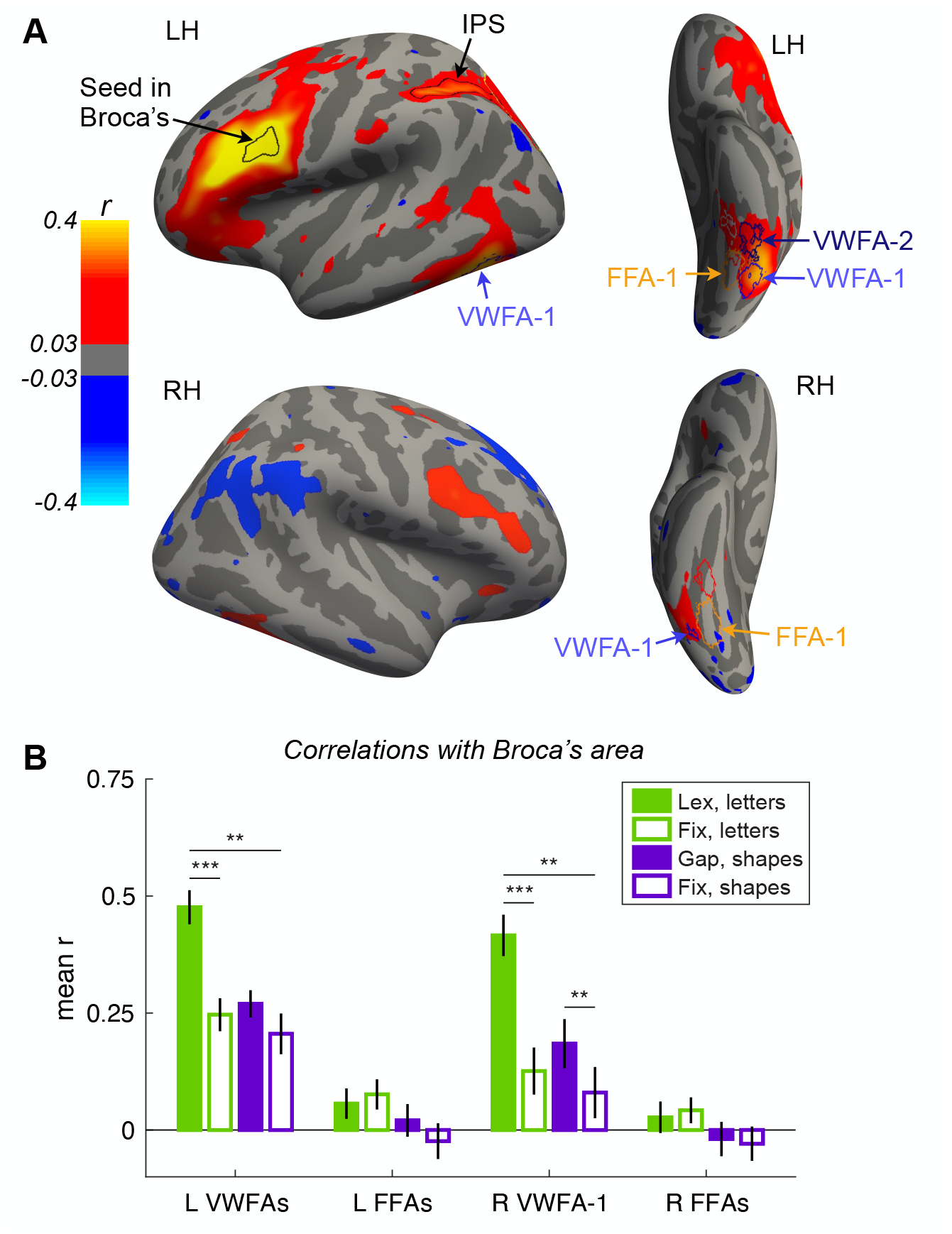
Functional connectivity with Broca’s area. **A**: Maps of mean correlations with Broca’s area during the lexical decision task, on Freesurfer’s “fsaverage” surface. The data are masked to show only vertices where the correlation was significant (corrected for false discovery rate < 0.05), peaking at *r*>=0.4. Lateral and ventral views of both hemispheres (LH and RH) are pictured. The “Broca’s area” ROI, defined from a univariate contrast of letters>shapes, is outlined in black, as is an IPS region drawn to encompass r>0.2. Just posterior to that, outlined in yellow, is IPS-0, -1 and -2. **B**: Mean correlation coefficients between Broca’s area and ventral temporal regions, in each condition. “VWFAs” is the average across VWFA-1 and -2, and “FFAs” is the average across FFA-1 and -2. Asterisks and abbreviations as in Figure 2. See also Figure S4.

In summary, the left VWFAs and Broca’s area showed a unique pattern: compared to when stimuli were ignored during the fixation task, responses to letters were greatly enhanced during the lexical task, but responses to shapes were suppressed during the gap task. That interaction is markedly different from the general attentional amplifications typically observed.

### The VWFAs are sensitive to lexicality and more so during the lexical task

The highly specific task effects suggest that, beyond encoding orthographic information, the VWFAs execute computations that are specifically related to lexical tasks. To go further, we separately analyzed responses to pseudowords and real words (**Figure 4)**. The VWFAs are known to respond more strongly to pseudowords, likely because pseudowords require more time or more effort to process ^11,15,25,26,50,51^. Here we investigated whether the magnitude of that “lexicality effect” differed across tasks and stimulus positions. Within each hemisphere we pooled data from VWFA-1 and VWFA-2 because their lexicality effects did not differ in either task (*Ps*>0.3, BFs<0.5). The same was true for the FFAs.

In the left VWFAs (Fig. 4, upper left) responses were larger for pseudowords than real words (F(1,649)=73.4, *P*<10^−16^; BF=10^8^). The pseudoword>real word lexicality effect was present in both tasks assessed separately, but significantly smaller in the fixation task (interaction F(1,649)=18.0, *P*<10^−4^; BF=78.6). In the lexical task, mean ratio of pseudoword response to real word response was 1.62, and the mean difference was 0.12 p.s.c (SEM=0.02, t(14)=6.1, *P*=3×10^−5^; BF=888). In the fixation task, the effect was less than half as large: the mean pseudo:real ratio was 1.28, and the mean difference only 0.04 p.s.c. (SEM=0.014; t(14)=2.55, *P*=0.023; BF=2.97).

Also, the lexicality effect decreased with absolute stimulus eccentricity (F(1,649)=35.7, *P*=10^−8^, BF=8332). That was true for both tasks separately (both P<0.019). In the lexical decision task, pseudowords evoked larger responses than real words only within 6º eccentricity. This corresponds to task accuracy, which was near chance beyond 6º (Fig. 1D). Thus, although mean BOLD responses during the lexical task were high for stimuli across the visual field (Fig. 2), the VWFA’s *differential* response to pseudowords vs. real words roughly mirrors participants’ ability to distinguish the stimuli.

We also analyzed lexicality effects in four other areas. The right VWFAs (Fig. 4, upper right) differentiated real and pseudowords much like the left VWFAs did. In the left FFAs (middle row left), there was a hint of a lexicality x eccentricity interaction during the lexical decision task, but the BF favored the null hypothesis. The right FFAs were clearly unaffected by lexicality. Responses in Broca’s area (bottom left) were more like the VWFAs: a strong effect of lexicality (pseudo>real; F(1,649)=27.6, *P*=10^−6^, BF=67.2), which interacted with task (F(1,649)=12.2, *P*=0.001, BF=3.3) and eccentricity (F(1,649)=15.2, *P*=10^−4^). Broca’s area was not affected by lexicality during the fixation task (F(1, 326)=1.56, *P*=0.21, BF=0.001).

Thus, the lexicality effect is specific to the VWFAs and Broca’s area and is much stronger when the participant engages in the lexical task. We propose that the response elevation to pseudowords is caused by prolonged voluntary effort to match the letter string to a known word. Familiar real words, in contrast, are rapidly matched and evoke a weaker BOLD response. That effort is not made when the word is so far in the periphery that it is completely illegible, nor during the fixation task (except perhaps on some trials when a word appears in the fovea and attention “leaks” to it from the fixation dot).

### Task-dependent connectivity between VWFA and Broca’s area

One explanation for the task effects shown above (Figs 2-4) is that voluntary effort to recognize words elicits top-down feedback from language areas, including Broca’s area, that specifically targets the left VWFAs. If so, incidental trial-to-trial fluctuations should be correlated between Broca’s area and the VWFAs. To test that prediction, we did a functional connectivity analysis with Broca’s area as the “seed.” For each surface vertex, we extracted the “residuals” in single-trial responses by subtracting out the across-trial mean response for each stimulus type (real words, pseudowords, shape strings), task, and stimulus position. Then we computed the correlation between each vertex’s residuals and the mean residuals in Broca’s area.

**Figure 5A** shows surface maps of those mean correlation coefficients during the lexical task. Broca’s area is a hotspot because it correlates well with itself. The correlations are high in the left VWFAs, but not in the neighboring face areas. Right VWFA-1 also shows significant correlation with Broca’s, as does the left intraparietal sulcus (IPS), and a right hemisphere homologue of Broca’s area. These patterns are specific: it is not just that the VWFAs correlate with many brain areas. **Supplementary Figure S4A** demonstrates that when left VWFA-1 is the “seed”, the same patches of cortex show significant correlation. For pairwise correlations between 14 different regions in each task condition, see **Figure S4B**.

More importantly, functional connectivity depended jointly on stimulus type and task. **Figure 5B** plots mean correlation coefficients with Broca’s area extracted from key ROIs. In the left VWFAs (mean of VWFA-1 and -2), the correlation was high (r=0.48) during the lexical decision task, but roughly half as strong in all other conditions. When letters were on the screen but ignored (fixation task), the correlation was not even as strong as when shapes were attended (gap task). The effect of task on the correlation for trials with letters was large (t(14) = 6.54, *P*=10^−5^, BF=1732), but there was little to no task effect for shapes (t(14) = 1.43, *P*=0.17, BF=0.61), and there was a strong interaction (t(14)=3.77, *P*=0.002, BF=21). Within the fixation task, the correlation when letters were presented was not any stronger than when shapes were presented (t(14)=1.29, *P*=0.22; BF=0.52).

When analyzed separately, VWFA-1 and VWFA-2 were similarly correlated with Broca’s (during the lexical task, r=0.52 vs. r=0.43; *P*=0.28, BF=0.47). Right VWFA-1 showed a similar correlation pattern as the left VWFAs, except with a significantly stronger correlation for shapes during the gap task than fixation task (t(10)=3.49, *P*=0.006; BF=9.4). The FFAs showed very little correlation with Broca’s, and no effects of task, although the left FFAs did have a slightly stronger correlation when letters were present than shapes (F(1,56)=7.40, *P*=0.009). For a full analysis of activity in the IPS region, see **Supplemental Figure S5**.

## DISCUSSION

By carefully manipulating both stimulus parameters and the participant’s task, we revealed highly specific modulations of BOLD activity in ventral occipito-temporal cortex. By “highly specific” we mean that within this data set, the connectivity patterns and task effects were restricted to text-selective regions. Instructing the participant to engage in a lexical decision task nearly doubled the VWFA’s responses to words across the visual field, compared to when the participant focused on the color of the fixation dot (Fig. 2). The difference in response between real words and novel pseudowords was also enhanced (Fig. 4; see also ref. ^15^). Remarkably, when the participant judged the location of a gap in a string of shapes, the VWFA’s response was *reduced* compared to when those shapes were ignored during the fixation task (Fig 2). This reversal of task-relevance effects was specific to the VWFAs and Broca’s area, and is the opposite of the expected enhancement for attended stimuli – which did arise in some retinotopic areas (Fig. 3). We are not aware of any other examples of visual stimuli evoking smaller fMRI responses when task-relevant than when ignored. This highlights the importance of modulations in visual cortex that are driven by task-specific cognitive processes.

For the first time, we combined task effects on mean BOLD responses with functional connectivity to better understand the network architecture. Trial-to-trial fluctuations were selectively correlated between the VWFAs, a putative left frontal Broca’s area, and the IPS. Compared to other visual areas, the VWFAs had privileged connectivity to Broca’s area in all conditions. However, that activity correlation was roughly twice as strong during the lexical task as the other tasks (Fig 5). That strong connectivity requires engagement in a lexical task, beyond the mere presence of words, because during the fixation task the correlation was not higher for words than shapes.

Altogether, these results suggest that the reading network is activated by voluntary engagement in a word recognition task, and can be suppressed by ignoring words or engaging in a non-linguistic task on word-like stimuli. One hypothesis we favor is that Broca’s area (perhaps together with the IPS) is a source of control for other parts of the reading network. We consider it as a source of “linguistic effort.” When Broca’s area is engaged, it communicates with the VWFA and boosts activity there, especially when a letter string is unfamiliar or difficult to recognize. These areas communicate iteratively until the letter string is either matched to an item in the mental lexicon or dismissed as meaningless. Thus, during the lexical task, the BOLD response in the VWFAs is elevated for pseudowords and for words in the parafovea. When the participant engages in the gap task, Broca’s area is less active and the VWFAs are suppressed. One possibility is that there is some mutual communication between Broca’s area and the VWFA even when non-linguistic stimuli are presented. During the gap task, a diversion of processing resources from the VWFA could lead to lower activity in Broca’s area, or vice versa, even if the strength of their correlation is unchanged.

In theory, the lexicality effect (pseudo>real words) could originate in Broca’s area and then be fed back to the VWFAs. However, that hypothesis is not supported by intracranial recordings: sensitivity to lexical features emerges early in a mid-fusiform text-selective area, not later than in frontal areas ^7^. An alternate hypothesis, therefore, is that orthographic lexical access first occurs locally in ventral temporal cortex, but doing so requires sustained excitatory feedback from frontal or parietal cortex.

One important question is whether such task-related modulations are unique to lexical tasks and to the VWFA, or whether analogous modulations occur for other tasks and in other category-selective regions. As one step towards answering this question, we analyzed data from a previous study^47^ in which three participants viewed sequences of face images presented at various positions and performed three different tasks: (1) Face identity task: attend to the faces to detect when two sequential images are of the same person (but viewed from different angles); (2) Face dot task: attend to the faces to detect a red dot occasionally superimposed on one of them; (3) Fixation digit task: ignore the faces to detect repetitions within numerals flashing at the screen center. Responses in the right FFAs were shown to be highest in the face identification task and lowest in the fixation task ^47^. In a new analysis, we found that this task effect on responses to faces was widespread in both hemispheres (**Figure 6)**. Even the left VWFAs responded more strongly in the face identification task than in the face dot task and the fixation task.

**Figure 6:**
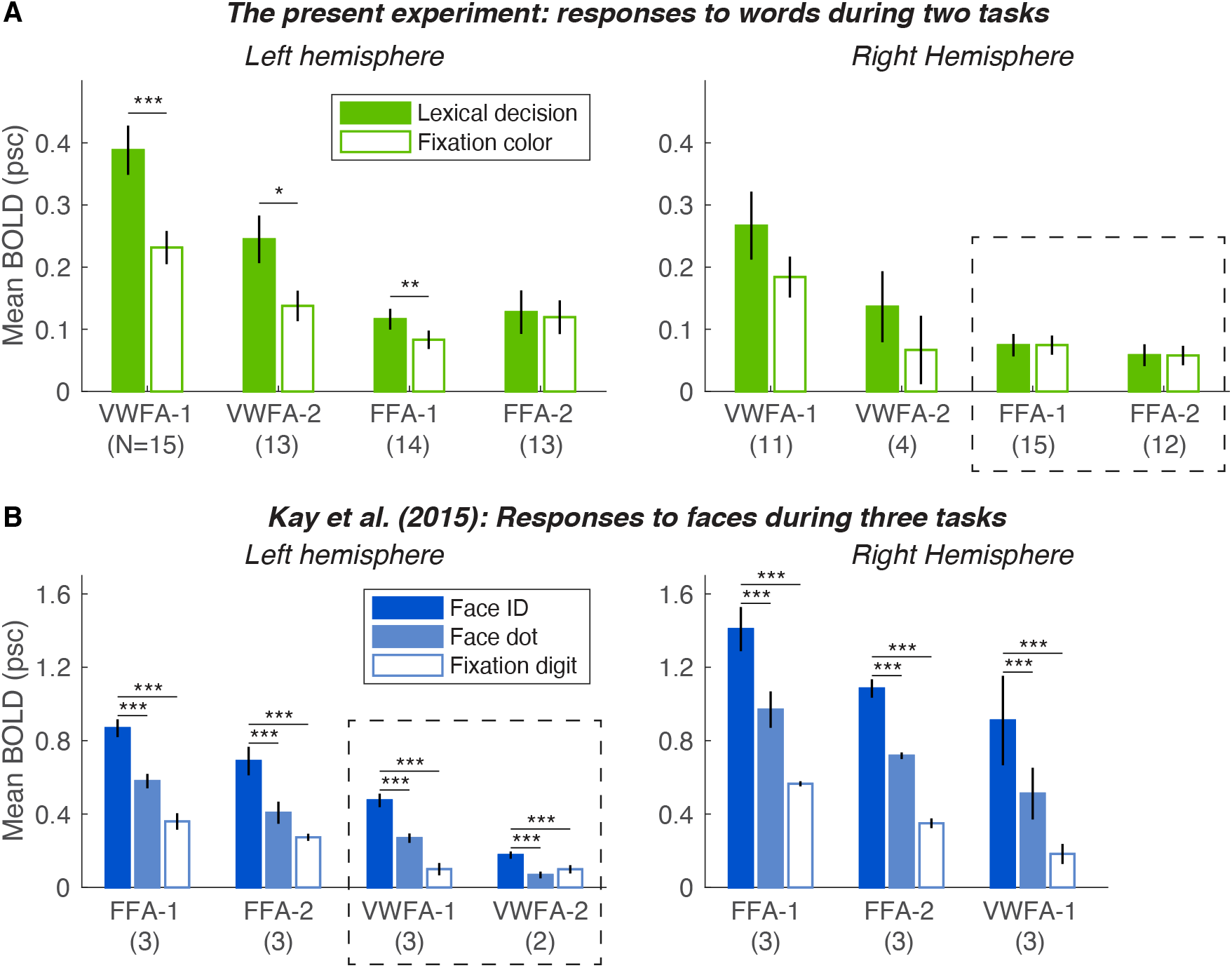
Less specific top-down effects during face recognition tasks. This figure compares responses to words in the current study (top row) with responses to faces in data from ref. ^47^ (bottom row). (**A**) The lexical>fixation task effect on responses to words was strong in the left VWFAs (left being the dominant hemisphere for word recognition) but absent in the right FFAs (dashed outline; right being the dominant hemisphere for faces). (**B**) Task effects on responses to faces were widespread in both hemispheres. Even in the left VWFAs (in dashed rectangle), responses were stronger in the face identity task than the dot task. In both those tasks, spatial attention was focused on the faces. Therefore, the effects on BOLD responses are due to more than just directing spatial attention to the stimuli. The parenthetical numbers below each ROI name indicate the number of participants. Asterisks as in Figure 2.

Therefore, engaging in a face identification task recruits swaths of visual cortex beyond the face-selective areas. Some caveats, however: several details of the stimulus presentation (e.g., 4 s vs 150 ms) differed between studies, and the face study^47^ collected a vast amount of data from only three participants. Nonetheless, the overall patterns are consistent with data from a larger sample, in which the VWFAs and FFAs responded more strongly to both faces and words in a one-back task than in a fixation task ^14^. In contrast, here we found that engaging in a word recognition task recruits the VWFAs specifically, due to top-down influences from language cortex that are spatially focused. This is evidence that the top-down effects we report above are a unique aspect of reading.

The suppression of VWFA activity we found in the gap task also stands apart from the face task data and other studies. It is *not* generally true that ventral temporal visual regions are suppressed during tasks that do not match their selectivity: in our data, the FFAs were not suppressed during either the lexical or gap tasks. Also, in the re-analyzed data with responses to face images ^47^, the VWFAs were not suppressed during the face tasks. We hypothesize that such a suppression may depend on stimuli that are *roughly* matched to a region’s preference, because our shape strings were visually similar to letter strings (unlike faces). This is an important open question for future research.

One common finding in our study and its predecessors is the involvement of the IPS. Kay & Yeatman (2017) concluded that areas IPS-0 and IPS-1 integrate sensory information from ventral temporal regions and send feedback that boosts weak signals ^14^, see also ^52,53^. These findings are consistent with IPS being part of a dorsal attentional control system ^54,55^. However, the part of the IPS highlighted by our functional connectivity analysis extends more anterior than the specific regions (e.g., IPS-0/1) associated with visual attention^14^ (see Figs. 5 and S4). The more anterior IPS is associated with letter position encoding ^56^, lexical processing ^7^, orthographic working memory ^57^, and even semantic memory ^58^. Other authors have also found IPS to be functionally connected to both the VWFA and Broca’s area ^38,42,59–61^. Therefore, anterior IPS may work with left frontal language regions to modulate visual word processing.

Another insight provided by our data is that the lexical task boosts responses to words even when they are presented in peripheral vision. Previous studies reported that the left VWFA has a limited “field of view” that extends only a few degrees to the left of fixation and drops off quickly between 5-10º to the right ^21,22^. Those studies measured responses to a sweeping bar during a fixation task. We found a flatter spatial profile during the lexical decision task: single flashed words evoked large responses even at 9º eccentricity to the left and right. However, the *difference* in response between words and pseudowords was more limited to the central visual field (Fig. 4), roughly matching behavioral accuracy. Therefore, the behaviorally relevant field of view – corresponding to how well the person can recognize words – may not be a simple reflection of the overall BOLD response.

One important question arising from this study is: are the task effects due to “visual attention”? In a loose sense, they must be, because the tasks required the participant to pay attention to different things. But we can be more specific. Covert, endogenous spatial attention probably plays a role: the fixation task required a narrow focus on just the very center of the screen, whereas both the lexical and gap tasks required an even distribution of attention across all possible stimulus locations. Indeed, in area hV4 we found a general enhancement of responses in the lexical and gap tasks. But spatial attention alone cannot explain why in the VWFA, responses to letters were enhanced, while responses to shapes were suppressed. We must invoke other mechanisms to explain our results. Another form of attention is feature-based: a boost in activity for neurons that are tuned to task-relevant features, like colors or motion directions ^5,62,63^. Stimuli that do not match a neuron’s preference can evoke smaller responses when attended than when totally ignored ^64^. Our tasks differed in the task-relevant features (red color, high spatial frequency contours, letter identities, etc.). Could some combination of feature-based and spatial attention explain our data? Perhaps, but only if we extend our concept of feature-based attention to include a dimension along which the features relevant for the lexical and gap tasks are at opposite ends, with the features relevant for the fixation task in the middle. The face task data further complicate this explanation because faces have very different features from words and yet evoke stronger VWFA responses when attended than ignored (Fig. S6).

Rather than drastically stretch the models of visual attention that have been elegantly applied to visual cortex in the past, we argue for a more inclusive view of task effects beyond attention. In other contexts, top-down modulations in ventral temporal cortex have proved more complex than spatial and feature-based attention ^52^. Our data implicate flexible and specific integration of visual and linguistic information that depends on top-down signals to support reading.

Going forward, a major goal is to develop a quantitative model of how activity in ventral temporal cortex depends on both stimulus features and task demands. Some conditions remain to be tested: for instance, how the VWFAs respond to non-linguistic stimuli during a lexical task, or to words during a non-linguistic task. Moreover, additional studies are necessary to determine whether the patterns observed here are unique to linguistic processing in the VWFAs, or whether analogous effects could arise elsewhere. Although existing data (Fig. S6) suggest that a face identification task evokes more wide-spread modulations, we must design tasks analogous to those used here but optimized for other brain areas. Such data, and a model that accounts for them, will further illuminate the extent to which cortical regions are distinct modules or components of integrated networks.

## ACKNOWLEDGEMENTS

We are grateful to Brian Wandell, Kalanit Grill-Spector, and Anthony Norcia for help designing the study, and to Vassiki Chauhan for writing advice. Funding provided by NIH R00 EY029366, R01 HD095861, and P41 EB027061; NSF IIS-1822683 and IIS-1822929. Thanks also to Kalinit Grill-Spector and Kevin Weiner for permission to re-analyze their 2015 data, funded by NIH R01 EY02391501A1 and RO1 EY03164.

## AUTHOR CONTRIBUTIONS

Conceptualization and Methodology, A.L.W, J.D.Y and K.K.; Software, A.L.W. and K.K., Investigation and Data Curation, K.A.T. and A.L.W., Formal Analysis, Visualization, and Writing – Original Draft, A.L.W., Writing – Reviewing & Editing, J.D.Y and K.K., Supervision, J.D.Y. and K.K., Resources, J.D.Y, Funding Acquisition, J.D.Y. and A.L.W.

## INCLUSION AND DIVERSITY

Two of the authors identify as members of the LGBTQ+ community. We support inclusive and equitable conduct of research.

## DECLARATION OF INTERESTS

The authors declare no competing interests.

## METHODS

### RESOURCE AVAILABILITY

#### Lead contact

Further information and requests for resources should be directed to and will be fulfilled by the lead contact, Alex White

#### Materials availability

This study did not generate new materials.

#### Data and code availability

- De-identified raw and pre-processed MRI data have been deposited at OpenNeuro: https://doi.org/10.18112/openneuro.ds004489.v1.0.0
- Processed data and original analysis code have been the Open Science Framework: https://doi.org/10.17605/OSF.IO/YU8SW
- Any additional information required to reanalyze the data reported in this paper is available from the lead contact upon request.

### EXPERIMENTAL MODEL AND SUBJECT DETAILS

This study was approved by the Institutional Review Board of Stanford University and complies with all relevant ethical regulations. All participants gave written informed consent and were paid a fixed monetary reward.

15 volunteers (10 female) participated. Their ages ranged from 19 to 28 years (mean = 23.8), and 14 were right-handed. All had normal or corrected to normal vision, and no history of dyslexia or other cognitive disorders. All scored at or above the population norm of 100 on the TOWRE-II tests of sight word efficiency and phonemic decoding efficiency ^65^: means (and SDs) = 120 (9) and 117 (9), respectively. One additional participant was excluded from the analyses for falling asleep in nearly every scan and performing near chance even for stimuli at fixation.

### METHOD DETAILS

#### Equipment

We acquired MRI data at the Center for Cognitive Neurobiological Imaging at Stanford University on a 3T GE Discovery MR750 scanner (GE Medical Systems) using a 32-channel head coil. In each session we collected one T1-weighted structural scan with 0.9 mm isotropic voxel size. We acquired functional data with a T2* sensitive gradient echo EPI sequence with a multiplexing (multiband) factor of 3 to acquire whole-brain coverage (51 slices). The TR was 1.19 s, TE was 30 ms and flip angle was 62°. The voxel size was 2.4 mm isotropic.

Via a mirror mounted above their nose, participants viewed the stimuli on an LCD screen (total viewing distance = 280 cm). The display had a resolution of 1920 × 1080 pixels, refreshing at 60 Hz. We presented the stimuli with custom MATLAB software (MathWorks, Natick, MA, USA) and the Psychophysics Toolbox ^66,67^. Throughout each scan we recorded monocular gaze position with an SR Research Eyelink 1000 tracker. Calibration was usually successful (details below), and even when it was not, participants believed their fixation was being monitored. Participants responded to the tasks by pressing two buttons on a response pad held in their right hand.

#### Main Experiment: Stimuli

**Figure 1A** shows two example stimuli. The screen’s background luminance was set to 80% of its maximum. The visual displays on each trial consisted of a persistent fixation dot of diameter 0.11 degrees of visual angle (dva) at the screen center, and one black stimulus string that flashed for 150 ms at a random one of 11 positions along the horizontal meridian. There were two stimulus types: letter strings and shape strings. The letter strings were all composed of 4 letters in “Liberation Mono” (a monospaced font similar to Courier). The font size was set such that the “x” was 0.41 dva tall. The 4-letter strings were on average 1.69 dva wide (range 1.62-1.76), with 0.44 degrees between the centers of neighboring letters. The stimulus set contained 264 unique letter strings, half of which were pronounceable pseudowords with constrained bigram statistics generated by MCWord ^68^. The other half were high-frequency real words of all syntactic categories (e.g., nouns and verbs). The mean frequency was 549 per million, ranging from 195 to 1,884. For the full list of stimuli, see **Supplemental Table 1**.

Each shape string was composed of 4 black circles and squares, matched in size and spacing to the letters (height = 0.43 dva, spacing = 0.44 dva). Each shape was composed of black lines 5 pixels wide (the same as most letter contours). There were 16 unique strings, each composed of 4 shapes that were independently and randomly set to be a square or a circle. Before being presented, one of the inner two letters had a gap added either to the top or the bottom side. This gap was 0.17 dva wide, equal in size to the gap in the letter “c”.

The fixation dot was usually dark gray in color (40% of maximum screen luminance), but on a random 50% of all trials it turned dark red during the 150 ms of stimulus presentation. When a stimulus was centered on fixation, the dot was superimposed onto the stimulus to remain visible.

#### Main experiment: Trials and tasks

Each 4-s trial was composed of one stimulus, a letter or shape string, presented for 150 ms, followed by a 3850 ms interval during which the subject could press a button to respond, followed immediately by the next trial. Trials came in blocks of 6. The stimulus type was constant within each block but varied randomly from block to block. Between blocks were blank periods of rest (no task except fixation on the dot), lasting 4, 6, or 8 seconds (randomly assigned).

The participant performed three different tasks at different times. Half of the runs (scans lasting ∼6 minutes) were “attend-fixation” and half were “attend-stimuli.” During “attend-fixation” runs, the participants ignored the letter and shape strings and performed the fixation task. The task was to press one of two keys to report whether or not the fixation dot turned red. The saturation of the red color (in HSV space) was controlled by a staircase to converge on the 80% correct detection threshold.

During the “attend-stimuli” runs, participants made judgments of the shape or letter strings while maintaining fixation on the dot but ignoring changes in its color. During blocks of trials with shape strings, the participant performed the gap task: to report whether the gap was on the top or bottom side. During blocks of trials with letter strings, the participant performed the lexical decision task: to report whether the presented letter string was a real word (e.g., book) or a pseudoword (e.g., blus). The task-irrelevant changes in fixation dot color were “replayed” from the staircases in fixation task runs.

Each subject saw each letter string once during the MRI experiment, and we took care to equalize the sets of stimuli presented at different locations in terms of metrics that could affect task difficulty and BOLD response. For each of the two run types (attend-stimuli, attend-fixation), we generated 11 sets of 12 letter strings (6 real and 6 pseudo-words), one set for each of the 11 visual field positions. Within each task, the word sets were balanced in terms of the average of four different metrics: log frequency (maximum spread = 0.1), orthographic neighborhood size (maximum spread = 3.5 for real words, 0.2 for pseudowords); mean RT from the English Lexicon Project (ELP) database ^69^ (maximum spread = 40 ms); and mean ELP accuracy (maximum spread = 5% correct). Having created 22 such lists of letter strings (two tasks x 11 positions), we then generated 5 unique word-to-position assignments by randomly shuffling those 22 lists five times. Each subject was given a random one of those 5 assignments. The use of each stimulus list in the lexical decision or fixation task was also randomized across subjects. It was therefore rare for any two subjects to see the same word at the same location in the same task.

#### Localizer Experiment

In order to localize word- and face-selective ROIs, participants completed a separate localizer experiment in their first scan session. Participants viewed sequences of images from 4 different categories: faces, objects, letter strings, and false fonts (Figure 1C). The letter strings were 5-letter real words, pseudowords, and consonant strings, each in two different fonts (Courier New and Sloan). For the false fonts, we used two different false characters matched in low-level visual features to the real fonts: a pseudo-Courier ^70^, and a pseudo-Sloan ^71^.

Each 4-second trial was composed of 4 frames presented in rapid sequence (700 ms each with 300 ms blanks between). Each frame contained 3 grayscale images: one small image at fixation (roughly 1.5 dva wide), and two large ones to either side (at 5.75 dva eccentricity, roughly 8 dva wide; see Fig 1A). A fixation dot 0.11 dva in diameter was superimposed on the center image. All frames on each trial contained images from the same category (e.g. 4 frames of faces).

The screen’s background was set to a medium gray (63% of its maximum white). Letters and false font characters were black. The face and object images were normalized such that pixel intensities spanned the full range [0 255], with mean equal to the background gray. The real words were all low in lexical frequency (<10 occurrences per million). The pseudowords were generated to be pronounceable with constrained bigram statistics ^68^.

The participant performed two different tasks in separate 5-minute scans: a fixation color task (press a button when the fixation dot changed color), and one-back task (press a button when the stimulus images repeat). Those “target events” (fixation color change or successive image repetition) occurred on a random 33% of trials (each set independently of the other). Each participant completed 4 localizer runs, alternating between the two tasks. To define ROIs, we analyzed localizer data from both tasks together.

#### Retinotopy experiment

Each participant also completed three 5-minute runs of a standard retinotopy experiment in which the participant fixated on a central spot while viewing a bar that moved across the visual field ^72^. The bar moved in 8 different directions, taking 32 s to cross the screen each time. The bar contained high-contrast patterns including faces and words, which changed 5 times per second. The participant’s task was to maintain central fixation and press a button whenever the fixation dot changed color.

To analyze these data, we used the analyzePRF toolbox to estimate the population receptive field (pRF) of each surface vertex ^73,74^. By visualizing the eccentricity and polar angle maps, we located borders between retinotopic areas ^75^. To analyze responses during the main experiment in these retinotopic areas, we selected subsets of voxels corresponding to each stimulus position. Each vertex was assigned to one of the positions if: the PRF model fit R^2^ > 0.2; the horizontal coordinate of its PRF center was within 1.5 dva; and distance of its vertical coordinate from the horizontal meridian was less than 1 SD of the PRF size. The response to each stimulus was then averaged only across vertices assigned to that stimulus position. Stimuli at +/-9 deg were excluded from analysis of retinotopic areas, because the displays used for retinotopic mapping did not extend out that far.

#### Procedure

Participants practiced the lexical decision and gap tasks for at least 2 one-hour sessions outside the scanner, with immediate feedback about gaze fixation as well as auditory feedback about accuracy on each trial. They then participated in two MRI scanning sessions. The goal was to complete 4 runs of the localizer and 3 runs of retinotopy in the first session, and 8 runs of the main experiment in the second.

The main experiment included 528 trials: 132 for each stimulus type (letters, shapes) and task (attend-stimuli, attend-fixation) combination. Within those, there were 12 trials per visual field position. In a few cases we collected 1 fewer run than planned due to time constraints. We excluded scans in which any framewise displacement due to head motion exceeded 2.4 mm (1 voxel). This applied to 4 scans from 1 subject and 3 scans from a second subject. When computing statistics, we weighted each participant’s data by the number of trials they completed.

### QUANTIFICATION AND STATISTICAL ANALYSIS

#### MRI data preprocessing

We used *fMRIPrep* 20.2.1 ^76,77^, which is based on *Nipype* 1.5.1 ^78,79^ to carry out the following pre-processing steps. T1-weighted (T1w) images were corrected for intensity non-uniformity (INU) with N4BiasFieldCorrection ^80^, distributed with ANTs 2.3.3 ^81^.

The T1w-reference was then skull-stripped with a *Nipype* implementation of the antsBrainExtraction.sh workflow (from ANTs), using OASIS30ANTs as target template. Brain tissue segmentation of cerebrospinal fluid (CSF), white-matter (WM) and gray-matter (GM) was performed on the brain-extracted T1w using fast (FSL 5.0.9) ^82^. A T1w-reference map was computed after registration of 2 T1w images (after INU-correction) using mri_robust_template (FreeSurfer 6.0.1) ^83^. Brain surfaces were reconstructed using recon-all (FreeSurfer 6.0.1) ^84^, and the brain mask estimated previously was refined with a custom variation of the method to reconcile ANTs-derived and FreeSurfer-derived segmentations of the cortical gray-matter of Mindboggle ^85^.

For each participant’s BOLD runs (across all tasks and sessions), the following preprocessing was performed: First, a reference volume and its skull-stripped version were generated by aligning a single-band reference scan. A B0-nonuniformity map (or *fieldmap*) was estimated based on two (or more) echo-planar imaging (EPI) references with opposing phase-encoding directions, with 3dQwarp (AFNI 20160207) ^86^. Based on the estimated susceptibility distortion, a corrected EPI (echo-planar imaging) reference was calculated for a more accurate co-registration with the anatomical reference. The BOLD reference was then co-registered to the T1w reference using bbregister (FreeSurfer) which implements boundary-based registration ^87^. Co-registration was configured with six degrees of freedom. Head-motion parameters with respect to the BOLD reference (transformation matrices, and six corresponding rotation and translation parameters) are estimated before any spatiotemporal filtering using mcflirt (FSL 5.0.9) ^88^. BOLD runs were slice-time corrected using 3dTshift from AFNI ^86^. Then, a reference volume and its skull-stripped version were generated using a custom methodology of *fMRIPrep*. The BOLD time-series (including slice-timing correction) were resampled onto their original, native space by applying a single, composite transform to correct for head-motion and susceptibility distortions. The BOLD time-series were also resampled onto the Freesurfer *fsnative* and *fsaverage* surfaces (the latter being a template of the average brain). Framewise displacement (FD) was computed for each functional run, using an implementation in *Nipype*, following Jenkinson ^88^ and Power ^89^. Gridded (volumetric) resamplings were performed using antsApplyTransforms (ANTs), configured with Lanczos interpolation to minimize the smoothing effects of other kernels ^90^. Non-gridded (surface) resamplings were performed using mri_vol2surf (FreeSurfer). Many internal operations of *fMRIPrep* use *Nilearn* 0.6.2, mostly within the functional processing workflow.

#### BOLD response estimation

For both the localizer and main experiment, we conducted GLMs to estimate BOLD responses to the stimuli on each trial. These responses (beta weights) are expressed in percent signal change, and reflect changes relative to the “blank” periods when the participant was simply fixating a dot on an otherwise empty screen. For the localizer data, we used GLMdenoise ^91^ to estimate the across-trial mean beta weight for each stimulus category. For the main experiment, we used GLMsingle ^92,93^ to estimate single-trial beta weights. The design matrix coded each 150 ms stimulus presentation as a separate event. Both GLMdenoise and GLMsingle optimize the assumed hemodynamic response functions and remove from the final estimations a set of noise regressors unrelated to the task and stimulus. Compared to GLMdenoise, GLMsingle estimates hemodynamic response functions on a per-voxel basis and also introduces ridge regression to improve stability and accuracy of single-trial beta weights.

#### ROI definition

We defined word- and face-selective regions of interests (ROIs) using data from the localizer experiment (**Figure 1C**). We computed contrasts of BOLD responses during the localizer experiment (text vs. false fonts, faces and objects; faces vs. text, false fonts and objects). We drew the ROIs based upon a visualization of the contrast t-statistic (t>3) on each subject’s native surface.

We defined three text-selective ROIs: (1) VWFA-1 is anterior to the V4/VO1 border, in the posterior occipito-temporal sulcus (OTS). (2) VWFA-2 is anterior to VWFA-1, usually in the OTS but sometimes extending onto the gyri on either side. In some subjects VWFA-2 and VWFA-1 appeared contiguous at the chosen contrast threshold, but they always had separate peaks of text selectivity. We based these regions upon a prior publication ^18^. (3) Finally, we noted in some participants (9/15) a third text-selective region medial to VWFA-2, near the mid-fusiform sulcus (MFS), which we termed text-mfs. The text-mfs region may be the same as reported in electrocorticography studies ^11,94^ and an fMRI study ^27^. Some participants also had text-selective blobs posterior to VWFA-1, but we did not analyze them as they were highly variable and often extended into the area occupied by hV4.

In addition, we defined two face-selective regions in the fusiform gyrus: one relatively posterior and usually medial to VWFA-1, which we call FFA-1, and another medial to VWFA-2, which we call FFA-2 (similar to what others have called pFus-faces and mFus-faces; ^12^). **Figure 1D** displays the most likely locations of each ROI on the fsaverage surfaces. For images of each participant’s ROIs, see the public data repository. The table below notes the number of participants who had each ROI in each hemisphere.

We also defined early visual areas (bilateral V1, V2, V3, hV4, and VO1/2) with data from separate retinotopic mapping scans ^95^. To analyze responses during the main experiment in these retinotopic areas, we selected subsets of voxels corresponding to each stimulus position. Stimuli at +/-9 deg were excluded from analysis of retinotopic areas, because the retinotopic mapping stimuli did not extend out that far. Although not clearly visible in our own retinotopy data, we also extracted the approximate locations of lateral occipital area LO-1/2 ^96^ based on a published atlas ^97^.

Finally, we defined a putative Broca’s area in the left pre-central sulcus, based on data in the main experiment. For each subject, at each surface vertex, we computed the mean difference in responses to letters vs. shapes, collapsing across positions and task conditions. Those difference maps were warped to the fsaverage template surface, and then we computed across-subject statistics at each vertex. We defined Broca’s area as a reliably text-selective blob where the t>5. We then back-transformed that ROI into each participant’s native surface.

#### Locations of and number of subjects with each ROI

In the table below, N is the number of participants in which we could localize each region, and *x, y, z* are mean coordinates in standard MNI152 space. (Broca’s area was defined based on across-participant average data, so N=15 by default).

| Region | Left Hemisphere |  |  |  | Right hemisphere |  |  |  |
| --- | --- | --- | --- | --- | --- | --- | --- | --- |
| | N | $x$ | $y$ | $z$ | N | $x$ | $y$ | $z$ |
| VWFA-1 | 15 | -43 | -61 | -10 | 11 | 46 | -58 | -11 |
| VWFA-2 | 13 | -46 | -45 | -18 | 5 | 48 | -48 | -17 |
| text-MFS | 9 | -36 | -40 | -21 | 4 | 38 | -42 | -22 |
| FFA-1 | 14 | -38 | -59 | -16 | 15 | 38 | -54 | -17 |
| FFA-2 | 13 | -39 | -42 | -20 | 12 | 41 | -37 | -21 |
| Broca's | (15) | -39 | 8 | 25 | - | - | - | - |

#### Statistical analyses

We used a combination of statistical tests to compare across-subject mean data to null hypotheses that predict no effect of some manipulation. In many cases we used linear mixed-effect models with random effects for participants. For models of the interaction of two or three predictor variables, the results are reported as repeated-measures ANOVAs (F-statistics and p-values). For tests of the effect of a mean difference between two conditions, we conducted paired t-tests. We also report Bayes Factors (BFs) for each test, to quantify strength of evidence. The BF is the ratio of the probability of the data under the alternate hypothesis (that two conditions differ), relative to the probability of the data under the null hypothesis ^98^. For example, a BF of 10 indicates that the data are ten times more likely under the alternate hypothesis than the null hypothesis. We computed BFs using the bayesFactor toolbox in MATLAB (https://github.com/klabhub/bayesFactor: DOI: 10.5281/zenodo.4394422).

#### Functional connectivity analysis

We assessed whole-cortex functional connectivity with two “seed” regions: left VWFA-1 and Broca’s area (see ROI definitions above). For each stimulus and task condition (e.g., real words during the lexical task), for each subject, for each surface vertex, we subtracted from each trial’s response the mean across all trials with that type of stimulus at the same location in the same task. Real words and pseudowords were treated separately in this analysis. Then, for each stimulus/task condition, we computed the correlation between each vertex’s de-meaned responses and the de-meaned responses in the seed area (averaged across its vertices). Each such correlation coefficient estimates the correlation in trial-to-trial variance not explained by overall effects of stimulus type, position, and task. To create the cortical surface maps of mean functional connectivity (Figs. 5 and S4), we first transformed each subject’s map to fsaverage space, smoothed them with a 2D Gaussian kernel (full-width at half-maximum = 5 mm), and then averaged across subjects.

## Notes

https://doi.org/10.17605/OSF.IO/YU8SW

https://doi.org/10.18112/openneuro.ds004489.v1.0.0

### Competing Interest Statement

The authors have declared no competing interest.

### Summary of Updates

Several sections updated and clarified, and a new Figure 6 to compare these results to a prior published dataset.

## REFERENCES

1. Roelfsema, P.R., and de Lange, F.P. (2016). Early Visual Cortex as a Multiscale Cognitive Blackboard. Annu. Rev. Vis. Sci. 2, 131–151. 10.1146/annurev-vision-111815-114443.

2. Maunsell, J.H.R. (2015). Neuronal Mechanisms of Visual Attention. Annu. Rev. Vis. Sci. 1, 373–391. 10.1146/annurev-vision-082114-035431.

3. Nobre, A.C., Kastner, S., and Beck, D.M. (2014). Neural Systems for Spatial Attention in the Human Brain 10.1093/oxfordhb/9780199675111.013.011.

4. Carrasco, M. (2011). Visual attention: The past 25 years. Vision Res. 51, 1484–1525. 10.1016/j.visres.2011.04.012.

5. Maunsell, J.H.R., and Treue, S. (2006). Feature-based attention in visual cortex. Trends Neurosci. 29, 317–322. 10.1016/j.tins.2006.04.001.

6. Yeatman, J.D., and White, A.L. (2021). Reading: The Confluence of Vision and Language. Annu. Rev. Vis. Sci. 7, 487–517. 10.1146/annurev-vision-093019-113509.

7. Woolnough, O., Donos, C., Curtis, A., Rollo, P.S., Roccaforte, Z.J., Dehaene, S., Fischer-Baum, S., and Tandon, N. (2022). A Spatiotemporal Map of Reading Aloud. J. Neurosci. 42, JN-RM-2324-21. 10.1523/jneurosci.2324-21.2022.

8. McCandliss, B.D., Cohen, L., and Dehaene, S. (2003). The visual word form area: Expertise for reading in the fusiform gyrus. Trends Cogn. Sci. 7, 293–299. 10.1016/S1364-6613(03)00134-7.

9. Caffarra, S., Karipidis, I.I., Yablonski, M., and Yeatman, J.D. (2021). Anatomy and physiology of word - selective visual cortex : from visual features to lexical processing. Brain Struct. Funct. 10.1007/s00429-021-02384-8.

10. Gaillard, R., Naccache, L., Pinel, P., Clémenceau, S., Volle, E., Hasboun, D., Dupont, S., Baulac, M., Dehaene, S., Adam, C., et al. (2006). Direct Intracranial, fMRI, and Lesion Evidence for the Causal Role of Left Inferotemporal Cortex in Reading. Neuron 50, 191–204. 10.1016/j.neuron.2006.03.031.

11. Woolnough, O., Donos, C., Rollo, P.S., Forseth, K.J., Lakretz, Y., Crone, N.E., Fischer-baum, S., Dehaene, S., and Tandon, N. (2021). Spatiotemporal dynamics of orthographic and lexical processing in the ventral visual pathway. Nat. Hum. Behav. 5, 389–398. 10.1038/s41562-020-00982-w.

12. Grill-Spector, K., and Weiner, K.S. (2014). The functional architecture of the ventral temporal cortex and its role in categorization. Nat. Rev. Neurosci. 15, 536–548. 10.1038/nrn3747.

13. Ben-Shachar, M., Dougherty, R.F., Deutsch, G.K., and Wandell, B.A. (2007). Differential sensitivity to words and shapes in ventral occipito-temporal cortex. Cereb. Cortex 17, 1604–1611. 10.1093/cercor/bhl071.

14. Kay, K.N., and Yeatman, J.D. (2017). Bottom-up and top-down computations in word- and face-selective cortex. Elife 6, 1–29. 10.7554/eLife.22341.

15. Mano, Q.R., Humphries, C., Desai, R.H., Seidenberg, M.S., Osmon, D.C., Stengel, B.C., and Binder, J.R. (2013). The role of left occipitotemporal cortex in reading: Reconciling stimulus, task, and lexicality effects. Cereb. Cortex 23, 988–1001. 10.1093/cercor/bhs093.

16. Lerma-Usabiaga, G., Carreiras, M., and Paz-Alonso, P.M. (2018). Converging evidence for functional and structural segregation within the left ventral occipitotemporal cortex in reading. Proc. Natl. Acad. Sci. 115, E9981–E9990. 10.1073/pnas.1803003115.

17. Vinckier, F., Dehaene, S., Jobert, A., Dubus, J.P., and Sigman, M. (2007). Hierarchical coding of letter strings in the ventral stream: Dissecting the inner organization of the visual word-form system. Neuron 55, 143–156. 10.1016/j.neuron.2007.05.031.

18. White, A.L., Palmer, J., Boynton, G.M., and Yeatman, J.D. (2019). Parallel spatial channels converge at a bottleneck in anterior word-selective cortex. Proc. Natl. Acad. Sci. 116, 10087–10096. 10.1073/pnas.1822137116.

19. Szwed, M., Dehaene, S., Kleinschmidt, A., Eger, E., Valabrègue, R., Amadon, A., and Cohen, L. (2011). Specialization for written words over objects in the visual cortex. Neuroimage 56, 330–344. 10.1016/j.neuroimage.2011.01.073.

20. Rauschecker, A.M., Bowen, R.F., Parvizi, J., and Wandell, B.A. (2012). Position sensitivity in the visual word form area. Proc. Natl. Acad. Sci. 109, E1568–E1577. 10.1073/pnas.1121304109.

21. Le, R.K., Witthoft, N., Ben-Shachar, M., and Wandell, B.A. (2017). The field of view available to the ventral occipito-temporal reading circuitry. J. Vis. 17, 1–19. 10.1167/17.4.6.

22. Lerma-usabiaga, G., Le, R., Gafni, C., Ben-shachar, M., and Wandell, B.A. (2021). Interpreting sensory and cognitive signals in the cortical reading network.

23. Hirshorn, E.A., Li, Y., Ward, M.J., Richardson, R.M., Fiez, J. a., and Ghuman, A.S. (2016). Decoding and disrupting left midfusiform gyrus activity during word reading. Proc. Natl. Acad. Sci. U. S. A. 113, 201604126. 10.1073/pnas.1604126113.

24. Glezer, L.S., Jiang, X., and Riesenhuber, M. (2009). Evidence for Highly Selective Neuronal Tuning to Whole Words in the “Visual Word Form Area.” Neuron 62, 199–204. 10.1016/j.neuron.2009.03.017.

25. Kronbichler, M., Bergmann, J., Hutzler, F., Staffen, W., Mair, A., Ladurner, G., and Wimmer, H. (2007). Taxi vs. taksi: On orthographic word recognition in the left ventral occipitotemporal cortex. J. Cogn. Neurosci. 19, 1584–1594. 10.1162/jocn.2007.19.10.1584.

26. Kronbichler, M., Hutzler, F., Wimmer, H., Mair, A., Staffen, W., and Ladurner, G. (2004). The visual word form area and the frequency with which words are encountered: Evidence from a parametric fMRI study. Neuroimage 21, 946–953. 10.1016/j.neuroimage.2003.10.021.

27. Bouhali, F., Bézagu, Z., Dehaene, S., and Cohen, L. (2019). A mesial-to-lateral dissociation for orthographic processing in the visual cortex. Proc. Natl. Acad. Sci. U. S. A. 116, 21936–21946. 10.1073/pnas.1904184116.

28. Reich, L., Szwed, M., Cohen, L., and Amedi, A. (2011). A ventral visual stream reading center independent of visual experience. Curr. Biol. 21, 363–368. 10.1016/j.cub.2011.01.040.

29. Dziegiel-Fivet, G., Plewko, J., Szczerbiński, M., Marchewka, A., Szwed, M., and Jednoróg, K. (2021). Neural network for Braille reading and the speech-reading convergence in the blind: Similarities and differences to visual reading. Neuroimage 231. 10.1016/j.neuroimage.2021.117851.

30. Siuda-Krzywicka, K., Bola, L., Paplinska, M., Sumera, E., Jednorog, K., Marchewka, A., Sliwinska, M.W., Amedi, A., and Szwed, M. (2016). Massive cortical reorganization in sighted braille readers. Elife 5, 1–26. 10.7554/eLife.10762.

31. Dehaene, S., Pegado, F., Braga, L.W., Ventura, P., Nunes Filho, G., Jobert, A., Dehaene-Lambertz, G., Kolinsky, R., Morais, J., and Cohen, L. (2010). How learning to read changes the cortical networks for vision and language. Science (80-.). 330, 1359–1364. 10.1126/science.1194140.

32. Ludersdorfer, P., Wimmer, H., Richlan, F., Schurz, M., Hutzler, F., and Kronbichler, M. (2016). Left ventral occipitotemporal activation during orthographic and semantic processing of auditory words. Neuroimage 124, 834–842. 10.1016/j.neuroimage.2015.09.039.

33. Planton, S., Chanoine, V., Sein, J., Anton, J.L., Nazarian, B., Pallier, C., and Pattamadilok, C. (2019). Top-down activation of the visuo-orthographic system during spoken sentence processing. Neuroimage 202. 10.1016/j.neuroimage.2019.116135.

34. Yoncheva, Y.N., Zevin, J.D., Maurer, U., and McCandliss, B.D. (2010). Auditory selective attention to speech modulates activity in the visual word form area. Cereb. Cortex 20, 622–632. 10.1093/cercor/bhp129.

35. O’Craven, K.M., and Kanwisher, N. (2000). Mental imagery of faces and places activates corresponding stimulus-specific brain regions. J. Cogn. Neurosci. 12, 1013–1023. 10.1162/08989290051137549.

36. Amedi, A., Jacobson, G., Hendler, T., Malach, R., and Zohary, E. (2002). Convergence of visual and tactile shape processing in the human lateral occipital complex zohary. Cereb. Cortex 12, 1202–1212. 10.1093/cercor/12.11.1202.

37. Price, C.J., and Devlin, J.T. (2011). The interactive account of ventral occipitotemporal contributions to reading. Trends Cogn Sci 15, 246–253. S1364-6613(11)00057-X [pii]10.1016/j.tics.2011.04.001.

38. Chen, L., Wassermann, D., Abrams, D.A., Kochalka, J., Gallardo-Diez, G., and Menon, V. (2019). The visual word form area (VWFA) is part of both language and attention circuitry. Nat. Commun. 10, 1–12. 10.1038/s41467-019-13634-z.

39. Yeatman, J.D., Rauschecker, A.M., and Wandell, B.A. (2013). Anatomy of the visual word form area: Adjacent cortical circuits and long-range white matter connections. Brain Lang. 125, 146–155. 10.1016/j.bandl.2012.04.010.

40. Bouhali, F., Thiebaut de Schotten, M., Pinel, P., Poupon, C., Mangin, J.-F., Dehaene, S., and Cohen, L. (2014). Anatomical Connections of the Visual Word Form Area. J. Neurosci. 34, 15402–15414. 10.1523/JNEUROSCI.4918-13.2014.

41. Saygin, Z.M., Osher, D.E., Norton, E.S., Youssoufian, D.A., Beach, S.D., Feather, J., Gaab, N., Gabrieli, J.D.E., and Kanwisher, N. (2016). Connectivity precedes function in the development of the visual word form area. Nat. Neurosci. 19, 1250–1255. 10.1038/nn.4354.

42. Vogel, A.C., Miezin, F.M., Petersen, S.E., and Schlaggar, B.L. (2012). The putative visual word form area is functionally connected to the dorsal attention network. Cereb. Cortex 22, 537–549. 10.1093/cercor/bhr100.

43. Stevens, W.D., Kravitz, D.J., Peng, C.S., Tessler, M.H., and Martin, A. (2017). Privileged functional connectivity between the visual word form area and the language system. J. Neurosci. 37, 5288–5297. 10.1523/JNEUROSCI.0138-17.2017.

44. Li, J., Osher, D.E., Hansen, H.A., and Saygin, Z.M. (2020). Innate connectivity patterns drive the development of the visual word form area. Sci. Rep., 1–12. 10.1038/s41598-020-75015-7.

45. Koyama, M.S., di Martino, A., Zuo, X.N., Kelly, C., Mennes, M., Jutagir, D.R., Castellanos, F.X., and Milham, M.P. (2011). Resting-state functional connectivity indexes reading competence in children and adults. J. Neurosci. 31, 8617–8624. 10.1523/JNEUROSCI.4865-10.2011.

46. López-Barroso, D., Thiebaut de Schotten, M., Morais, J., Kolinsky, R., Braga, L.W., Guerreiro-Tauil, A., Dehaene, S., and Cohen, L. (2020). Impact of literacy on the functional connectivity of vision and language related networks. Neuroimage 213. 10.1016/j.neuroimage.2020.116722.

47. Kay, K.N., Weiner, K.S.S., and Grill-Spector, K. (2015). Attention reduces spatial uncertainty in human ventral temporal cortex. Curr. Biol. 25, 595–600. 10.1016/j.cub.2014.12.050.

48. Binder, J.R., McKiernan, K.A., Parsons, M.E., Westbury, C.F., Possing, E.T., Kaufman, J.N., and Buchanan, L. (2003). Neural correlates of lexical access during visual word recognition. J. Cogn. Neurosci. 15, 372–393. 10.1162/089892903321593108.

49. Heim, S., Eickhoff, S.B., Ischebeck, A.K., Supp, G., and Amunts, K. (2007). Modality-independent involvement of the left BA 44 during lexical decision making. Brain Struct. Funct. 212, 95–106. 10.1007/s00429-007-0140-6.

50. Taylor, J.S.H., Rastle, K., and Davis, M.H. (2013). Can cognitive models explain brain activation during word and pseudoword reading? A meta-analysis of 36 neuroimaging studies. Psychol. Bull. 139, 766–791. 10.1037/a0030266.

51. Gagl, B., Richlan, F., Ludersdorfer, P., Sassenhagen, J., Eisenhauer, S., Gregorova, K., and Fiebach, C.J. (2022). The lexical categorization model: A computational model of left ventral occipitotemporal cortex activation in visual word recognition. PLoS Comput. Biol. 18, 1–33. 10.1371/journal.pcbi.1009995.

52. Zhang, R.Y., and Kay, K.N. (2020). Flexible top-down modulation in human ventral temporal cortex. Neuroimage 218, 116964. 10.1016/j.neuroimage.2020.116964.

53. Cohen, L., Dehaene, S., Vinckier, F., Jobert, A., and Montavont, A. (2008). Reading normal and degraded words: Contribution of the dorsal and ventral visual pathways. Neuroimage 40, 353–366. 10.1016/j.neuroimage.2007.11.036.

54. Lauritzen, T.Z., Esposito, M.D., Heeger, D.J., Silver, M. a, Wills, H., and Jr, H.H.W. (2009). Top-down flow of visual spatial attention signals from parietal to occipital cortex. J. Vis. 9(13):18, 1–14. 10.1167/9.13.18.Introduction.

55. Silver, M. a, Ress, D., Heeger, D.J., Michael, a, and Topographic, D.J.H. (2005). Topographic Maps of Visual Spatial Attention in Human Parietal Cortex. J. Neurophysiol. 94, 1358–1371. 10.1152/jn.01316.2004.

56. Ossmy, O., Ben-Shachar, M., and Mukamel, R. (2014). Decoding letter position in word reading. Cortex 59, 74–83. 10.1016/j.cortex.2014.07.002.

57. Rapp, B., Purcell, J., Hillis, A.E., Capasso, R., and Miceli, G. (2016). Neural bases of orthographic long-term memory and working memory in dysgraphia. Brain 139, 588–604. 10.1093/brain/awv348.

58. Forseth, K.J., Kadipasaoglu, C.M., Conner, C.R., Hickok, G., Knight, R.T., and Tandon, N. (2018). A lexical semantic hub for heteromodal naming in middle fusiform gyrus. Brain 141, 2112–2126. 10.1093/brain/awy120.

59. Wang, X., Caramazza, A., Peelen, M. V., Han, Z., and Bi, Y. (2015). Reading without speech sounds: VWFA and its connectivity in the congenitally deaf. Cereb. Cortex 25, 2416–2426. 10.1093/cercor/bhu044.

60. López-Barroso, D., Thiebaut de Schotten, M., Morais, J., Kolinsky, R., Braga, L.W., Guerreiro-Tauil, A., Dehaene, S., and Cohen, L. (2020). Impact of literacy on the functional connectivity of vision and language related networks. Neuroimage 213. 10.1016/j.neuroimage.2020.116722.

61. Makuuchi, M., and Friederici, A.D. (2013). Hierarchical functional connectivity between the core language system and the working memory system. Cortex 49, 2416–2423. 10.1016/j.cortex.2013.01.007.

62. White, A.L., Rolfs, M., and Carrasco, M. (2015). Stimulus competition mediates the joint effects of spatial and feature-based attention. J. Vis. 15, 1–21. https://doi.org/10.1167/15.14.7.

63. Liu, T. (2019). Feature-based attention: effects and control. Curr. Opin. Psychol. 29, 187–192. 10.1016/j.copsyc.2019.03.013.

64. Treue, S., and Martínez Trujillo, J.C. (1999). Feature-based attention influences motion processing gain in macaque visual cortex. Nature 399, 575–579. 10.1038/21176.

65. Torgesen, J., Rashotte, C., and Wagner, R. (1999). TOWRE-2: Test of Word Reading Efficiency, 2nd Ed. Austin, TX Pro-Ed.

66. Brainard, D.H. (1997). The psychophysics toolbox. Spat. Vis. 10, 443–446.

67. Pelli, D.G. (1997). The VideoToolbox software for visual psychophysics: Transforming numbers into movies. Spat. Vis. 10, 437–442.

68. Medler, D.A., and Binder, J.R. (2005). MCWord: An on-Line orthographic database of the English language. http://www.neuro.mcw.edu/mcword/.

69. Balota, D.A., Yap, M.J., Cortese, M.J., Hutchison, K. a, Kessler, B., Loftis, B., Neely, J.H., Nelson, D.L., Simpson, G.B., and Treiman, R. (2007). The English Lexicon Project. Behav. Res. Methods 39, 445–459. 10.3758/BF03193014.

70. Vidal, C., Content, A., and Chetail, F. (2017). BACS: The Brussels Artificial Character Sets for studies in cognitive psychology and neuroscience. Behav. Res. Methods 49, 2093–2112. 10.3758/s13428-016-0844-8.

71. Vildavski, V., Lo Verde, L., Blumberg, G., Parsey, J., and Norcia, A. (2021). PseudoSloan: A perimetric-complexity and area-controlled font for vision and reading research. J. Vis. 21, 2857. 10.1167/jov.21.9.2857.

72. Benson, N.C., Jamison, K.W., Arcaro, M.J., Vu, A.T., Glasser, M.F., Coalson, T.S., Van Essen, D.C., Yacoub, E., Ugurbil, K., Winawer, J., et al. (2018). The Human Connectome Project 7 Tesla retinotopy dataset: Description and population receptive field analysis. J. Vis. 18, 1–22. 10.1167/18.13.23.

73. Dumoulin, S.O., and Wandell, B.A. (2008). Population receptive field estimates in human visual cortex. Neuroimage 39, 647–660. 10.1016/j.neuroimage.2007.09.034.

74. Kay, K.N., Winawer, J., Mezer, A., and Wandell, B.A. (2013). Compressive spatial summation in human visual cortex. J. Neurophysiol. 110, 481–494. 10.1152/jn.00105.2013.

75. Winawer, J., and Witthoft, N. (2017). Identification of the ventral occipital visual field maps in the human brain. F1000Research 6, 1–18. 10.12688/f1000research.12364.1.

76. Esteban, O., Blair, R., Markiewicz, C.J., Berleant, S.L., Moodie, C., Ma, F., Isik, A.I., and Poldrack, R.A. (2018). FMRIPrep. Software Zenodo, 852659. https://doi.org/10.5281/zenodo.852659.

77. Esteban, O., Markiewicz, C.J., Blair, R.W., Moodie, C.A., Isik, A.I., Erramuzpe, A., Kent, J.D., Goncalves, M., DuPre, E., Snyder, M., et al. (2019). fMRIPrep: a robust preprocessing pipeline for functional MRI. Nat. Methods 16, 111–116. 10.1038/s41592-018-0235-4.

78. Gorgolewski, K.J., Krzysztof, K.J., Esteban, O., Markiewicz, C.J., Ziegler, E., Gage, D., Notter, M.P., and Jarecka, D. (2018). Nipype. Software Zenodo. https://doi.org/10.5281/zenodo.596855.

79. Gorgolewski, K., Burns, C.D., Madison, C., Clark, D., Halchenko, Y.O., Waskom, M.L., and Ghosh, S.S. (2011). Nipype: A flexible, lightweight and extensible neuroimaging data processing framework in Python. Front. Neuroinform. 5. 10.3389/fninf.2011.00013.

80. Tustison, N.J., Avants, B.B., Cook, P.A., Zheng, Y., Egan, A., Yushkevich, P.A., and Gee, J.C. (2010). N4ITK: Improved N3 bias correction. IEEE Trans. Med. Imaging 29, 1310–1320. 10.1109/TMI.2010.2046908.

81. Avants, B.B., Epstein, C.L., Grossman, M., and Gee, J.C. (2008). Symmetric diffeomorphic image registration with cross-correlation: Evaluating automated labeling of elderly and neurodegenerative brain. Med. Image Anal. 12, 26–41. 10.1016/j.media.2007.06.004.

82. Zhang, Y., Brady, M., and Smith, S. (2001). Segmentation of brain MR images through a hidden Markov random field model and the expectation-maximization algorithm. IEEE Trans. Med. Imaging 20, 45–57. 10.1109/42.906424.

83. Reuter, M., Rosas, H.D., and Fischl, B. (2010). Highly accurate inverse consistent registration: A robust approach. Neuroimage 53, 1181–1196. 10.1016/j.neuroimage.2010.07.020.

84. Dale, A.M., Fischl, B., and Sereno, M.I. (1999). Cortical Surface-Based Analysis. Neuroimage 9, 179–194. 10.1006/nimg.1998.0395.

85. Klein, A., Ghosh, S.S., Bao, F.S., Giard, J., Häme, Y., Stavsky, E., Lee, N., Rossa, B., Reuter, M., Chaibub Neto, E., et al. (2017). Mindboggling morphometry of human brains 10.1371/journal.pcbi.1005350.

86. Cox, R.W., and Hyde, J.S. (1997). Software Tools for Analysis and Visualization of FMRI Data NMR in Biomedicine, in press. NMR Biomed 10, 171–178.

87. Greve, D.N., and Fischl, B. (2009). Accurate and robust brain image alignment using boundary-based registration. Neuroimage 48, 63–72. 10.1016/j.neuroimage.2009.06.060.

88. Jenkinson, M., Bannister, P., Brady, M., and Smith, S. (2002). Improved Optimization for the Robust and Accurate Linear Registration and Motion Correction of Brain Images. Neuroimage 17, 825–841. 10.1006/nimg.2002.1132.

89. Power, J.D., Mitra, A., Laumann, T.O., Snyder, A.Z., Schlaggar, B.L., and Petersen, S.E. (2014). Methods to detect, characterize, and remove motion artifact in resting state fMRI. Neuroimage 84, 320–341. 10.1016/j.neuroimage.2013.08.048.

90. Lanczos, C. (1964). Evaluation of Noisy Data. J. Soc. Ind. Appl. Math. Ser. B Numer. Anal. 1, 76–85. 10.1137/0701007.

91. Kay, K.N., Rokem, A., Winawer, J., Dougherty, R.F., and Wandell, B.A. (2013). GLMdenoise: A fast, automated technique for denoising task-based fMRI data. Front. Neurosci. 7, 247. 10.3389/fnins.2013.00247.

92. Allen, E.J., St-Yves, G., Wu, Y., Breedlove, J.L., Prince, J.S., Dowdle, L.T., Nau, M., Caron, B., Pestilli, F., Charest, I., et al. (2022). A massive 7T fMRI dataset to bridge cognitive neuroscience and artificial intelligence. Nat. Neurosci. 25, 116–126. 10.1038/s41593-021-00962-x.

93. Prince, J.S., Charest, I., Kurzawski, J.W., Pyles, J.A., Tarr, M.J., and Kay, K.N. (2022). GLMsingle: a toolbox for improving single-trial fMRI response estimates. bioRxiv. https://doi.org/10.1101/2022.01.31.478431;

94. Lochy, A., Jacques, C., Maillard, L., Colnat-Coulbois, S., Rossion, B., and Jonas, J. (2018). Selective visual representation of letters and words in the left ventral occipito-temporal cortex with intracerebral recordings. Proc. Natl. Acad. Sci. 115, E7595–E7604. 10.1073/pnas.1718987115.

95. Benson, N.C., Jamison, K.W., Arcaro, M.J., Vu, A.T., Glasser, M.F., Coalson, T.S., Van Essen, D.C., Yacoub, E., Ugurbil, K., Winawer, J., et al. (2018). The Human Connectome Project 7 Tesla retinotopy dataset: Description and population receptive field analysis. J. Vis. 18, 1–22. 10.1167/18.13.23.

96. Larsson, J., and Heeger, D.J. (2006). Two Retinotopic Visual Areas in Human Lateral Occipital Cortex. J. Neurosci. 26, 13128–13142. 10.1523/JNEUROSCI.1657-06.2006.

97. Wang, L., Mruczek, R.E.B., Arcaro, M.J., and Kastner, S. (2015). Probabilistic maps of visual topography in human cortex. Cereb. Cortex 25, 3911–3931. 10.1093/cercor/bhu277.

98. Rouder, J.N., Speckman, P.L., Sun, D., Morey, R.D., and Iverson, G. (2009). Bayesian t tests for accepting and rejecting the null hypothesis. Psychon. Bull. Rev. 16, 225–237. 10.3758/PBR.16.2.225.

