## Supplemental text and figures for "Engaging in word recognition elicits highly specific modulations in visual cortex"

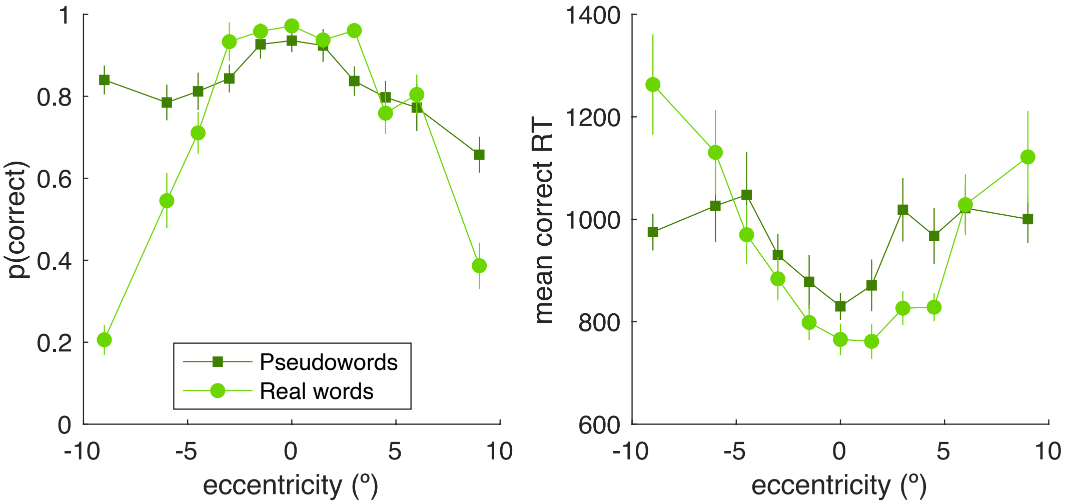


**Figure S1: Lexical decision task performance separately for real words and pseudowords, related to Figure 1B.** The left panel is mean accuracy (proportion trials correct), and the right panel is mean correct response time in ms. Error bars are ± 1 SEM across subjects. Accuracy drops off with eccentricity much more rapidly for real words (light green circles) than for pseudowords (dark green squares). This is consistent with a general bias participants have to report “pseudo” unless they clearly recognize the word. As the word visibility degrades with increasing eccentricity, reports of “real” become less and less common. Correct response times also tend to increase with eccentricity, for both types of stimuli.

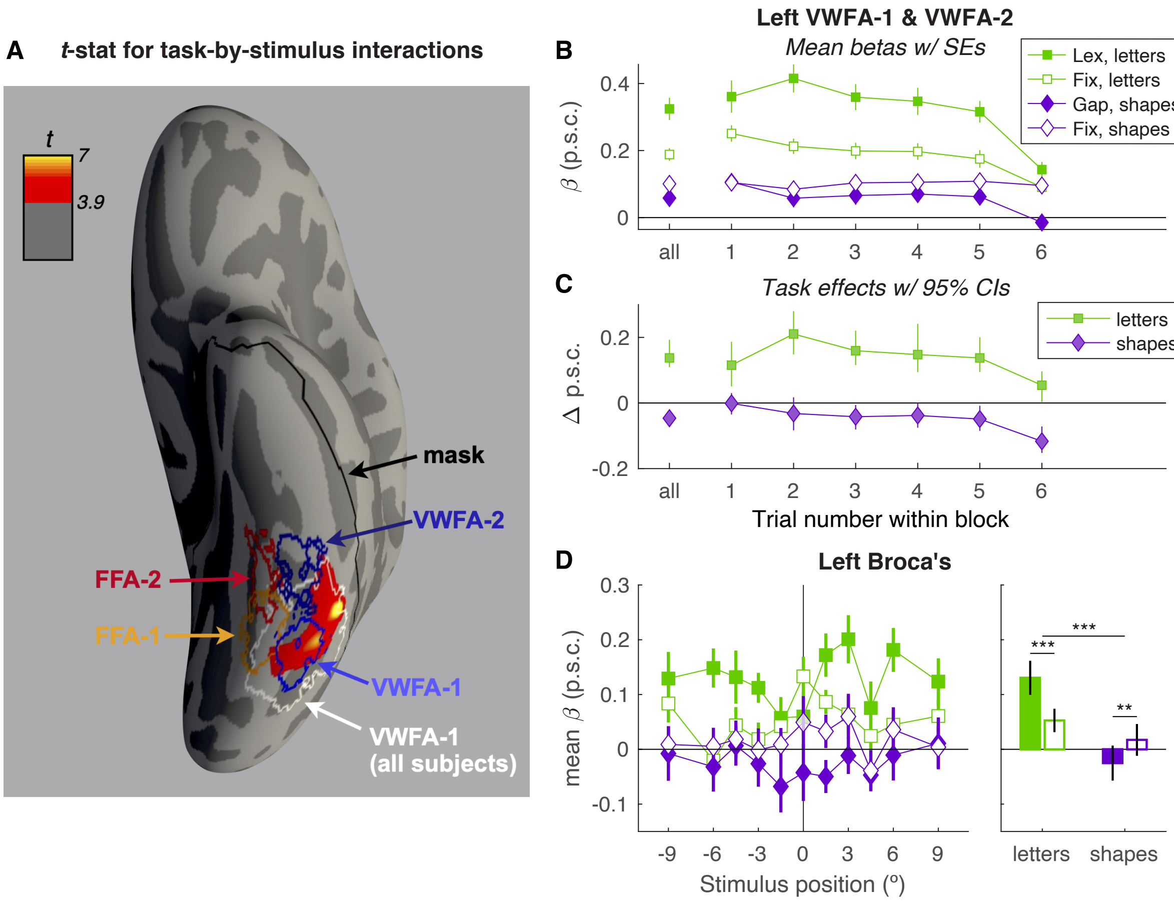


**Figure S2: specificity of the task-by-stimulus interaction, activity as a function of trial number, and activity in Left Broca's area. Related to Figures 2 and 3. A:** Left ventral cortical surface (fsaverage) with a map of the t-statistic for interaction in BOLD responses: (Lexical task – Fixation $task_{letters}$) – (Gap task – Fixation $task_{shapes}$). The colormap ranges from $3.9 \leq t \leq 7$, 3.9 being the minimum that was significant after correcting for multiple comparisons. The right hemisphere had no significant vertices. We analyzed these data only within the masked region, outlined in black, that included all of ventral temporal and early visual cortex (the union of Freesurfer parcellations fusiform, inferior temporal, parahippocampal, entorhinal, lateral occipital, lingual, pericalcarine, and cuneus.) "VWFA-1 (all subjects)" encompasses in white *all* individual VWFA-1 ROIs. The other ROIs are the same as those in main text Figure 1D. **B:** Mean responses as a function of trial number within the average block, in the union of left VWFA-1 and VWFA-2, collapsed across stimulus position. Trials came in blocks of 6 of all the same stimulus type. The stimulus type varied randomly from block to block. (The "all" condition on the left collapses across trial number within the block). Lex = lexical task, Gap=gap task, Fix=fixation task. Error bars are ± 1 SEM. Curiously, there was an overall trend for the absolute beta weights to decrease throughout the average block (slope = -0.02 psc per trial, $p<10^{-14}$). That could be due to repetition suppression or adaptation to repeated stimulation. **C:** The mean task effects as a function of trial number, in the left VWFAs. Each point is the difference in the corresponding two points in panel B, with 95% bootstrapped CIs. The positive task effect for letters is significant throughout, but there was a significantly negative linear effect of trial number (p=0.02, 95% CI of slope = [-0.03 -0.002]). The task effect for shapes was near 0 on the 1st trial, then became more negative (p=0.001, 95% CI of slope = [-0.03 -0.01]). **D:** Mean BOLD responses in the Broca's area ROI, analogous to main text Figure 2. Note that the task effect for letters (lexical>fixation enhancement) is smallest, or even reversed, for stimuli at the fovea (0°), but larger in the periphery. In contrast, the task effect for shapes (gap<fixation suppression) is largest near the fovea, where shapes actually induce a drop in the BOLD signal below baseline.

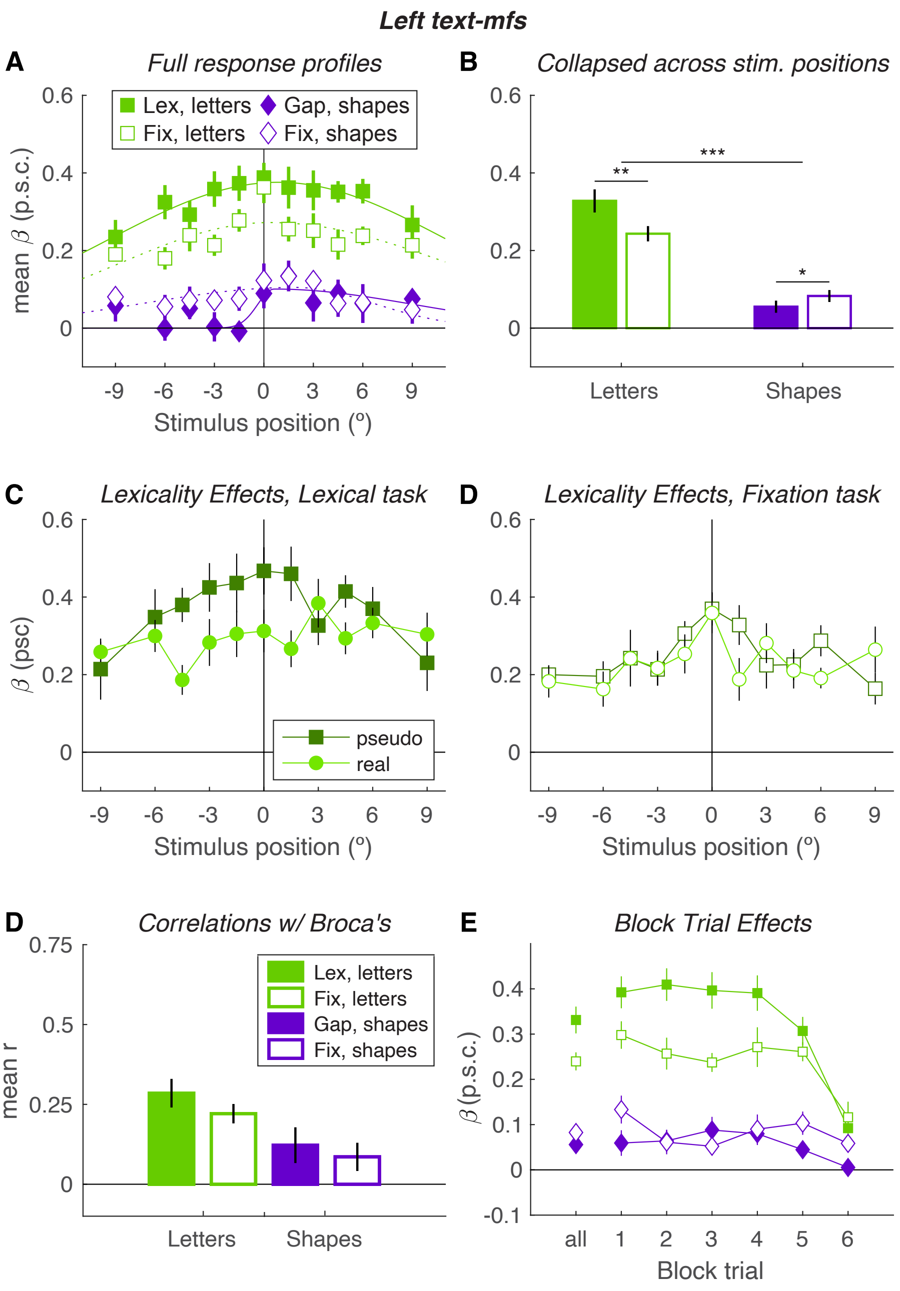


**Figure S3: Activity in the left text-mfs region, related to Figures 2, 3 and 4.** We could localize this region in only 9/15 participants. Each panel corresponds to a plot of VWFA activity in the main text. **A:** According an LME model, there was a significant main effect of absolute eccentricity ($p<10^{-6}$), which was larger for letters than shapes (p=0.007). **B:** as indicated by the asterisks, there were task effects for each stimulus type, which went in opposite directions. **C:** Responses were overall higher for pseudo than real words (F(1,210)=11, p=0.001), and more so in the lexical task (F(1,210)=3.9, p=0.049). **D:** The trial-to-trial response variability was more strongly correlated with Broca's area when letters were presented than shapes (p=0.008), but was not significantly affected by task. **E**: An LME model of response magnitudes revealed a main negative effect of trial (F(1, 208)=46, $p=10^{-10}$). The task effect for letters (lexical>fixation) significantly diminished across trials (t(52)=3.63, p=0.001), but the task effect for shapes was not affected (t<1, p>0.5). Unlike in the VWFAs, the gap<fixation effect was present on trial 1.

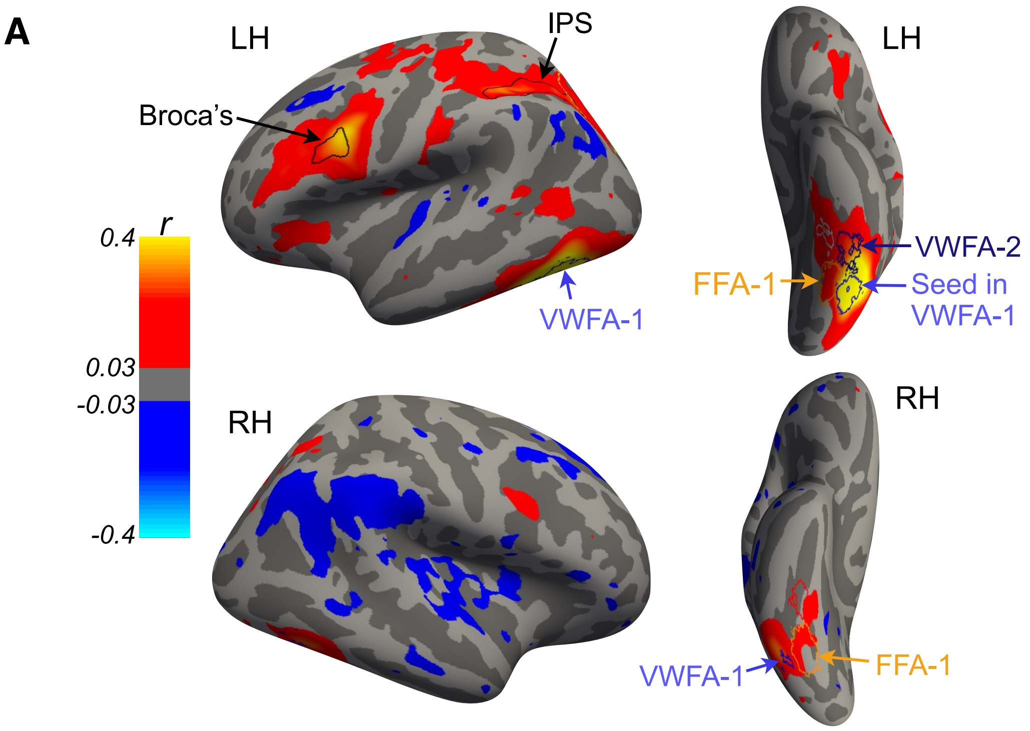


**B**

**Lexical Decision Task**

| | L V1 | L V4 | L LO | L VWFA-1 | L VWFA-2 | L text-mfs | L FFA-1 | L FFA-2 | R V1 | R VWFA-1 | R FFA-1 | R FFA-2 | L IPS |
|---|---|---|---|---|---|---|---|---|---|---|---|---|---|
| L V1 | | | | | | | | | | | | | |
| L V4 | 0.18* | | | | | | | | | | | | |
| L LO | 0.01 | 0.20** | | | | | | | | | | | |
| L VWFA-1 | 0.08 | 0.14 | 0.06 | | | | | | | | | | |
| L VWFA-2 | 0.06 | 0.04 | -0.08 | 0.56** | | | | | | | | | |
| L text-mfs | 0.19* | 0.13 | -0.04 | 0.54** | 0.30 | | | | | | | | |
| L FFA-1 | 0.08 | 0.30** | 0.24** | 0.32** | 0.17 | 0.30* | | | | | | | |
| L FFA-2 | 0.05 | 0.15 | 0.04 | 0.32** | 0.34* | 0.39** | 0.33** | | | | | | |
| R V1 | 0.45** | 0.32** | 0.17 | -0.07 | -0.01 | -0.08 | -0.07 | -0.05 | | | | | |
| R VWFA-1 | -0.02 | -0.04 | 0.05 | 0.55** | 0.39** | 0.29 | 0.18 | 0.16 | 0.01 | | | | |
| R FFA-1 | 0.03 | 0.16 | 0.12 | 0.13* | 0.10 | 0.15 | 0.36** | 0.24* | -0.05 | 0.29** | | | |
| R FFA-2 | 0.01 | 0.04 | 0.04 | 0.16 | 0.18 | 0.18 | 0.26* | 0.23* | -0.00 | 0.25 | 0.36* | | |
| L IPS | -0.03 | -0.02 | -0.01 | 0.50** | 0.38** | 0.25 | 0.04 | 0.15 | -0.08 | 0.43** | -0.01 | 0.01 | |
| L Broca's | -0.04 | -0.09 | -0.13 | 0.52** | 0.43** | 0.29** | 0.04 | 0.08 | -0.12* | 0.42** | -0.01 | 0.07 | 0.53** |

**Gap task**

| | L V1 | L V4 | L LO | L VWFA-1 | L VWFA-2 | L text-mfs | L FFA-1 | L FFA-2 | R V1 | R VWFA-1 | R FFA-1 | R FFA-2 | L IPS |
|---|---|---|---|---|---|---|---|---|---|---|---|---|---|
| L V1 | | | | | | | | | | | | | |
| L V4 | 0.22** | | | | | | | | | | | | |
| L LO | 0.14 | 0.29** | | | | | | | | | | | |
| L VWFA-1 | 0.07 | 0.13 | 0.09 | | | | | | | | | | |
| L VWFA-2 | -0.01 | 0.08* | -0.02 | 0.47** | | | | | | | | | |
| L text-mfs | 0.08 | 0.15 | 0.02 | 0.43** | 0.34** | | | | | | | | |
| L FFA-1 | 0.14 | 0.27** | 0.23* | 0.30** | 0.22** | 0.30** | | | | | | | |
| L FFA-2 | 0.08 | 0.08 | 0.04 | 0.28** | 0.34** | 0.30* | 0.29** | | | | | | |
| R V1 | 0.41 | 0.23 | -0.02 | -0.03 | -0.01 | 0.04 | 0.08 | 0.01 | | | | | |
| R VWFA-1 | 0.05 | 0.04 | 0.19* | 0.48** | 0.30** | 0.11 | 0.20* | 0.14* | 0.11 | | | | |
| R FFA-1 | 0.06 | 0.14 | 0.14 | 0.17** | 0.15* | 0.11 | 0.40** | 0.23* | 0.09 | 0.32** | | | |
| R FFA-2 | -0.02 | 0.02 | 0.04 | 0.20** | 0.20 | 0.15 | 0.18 | 0.21 | -0.05 | 0.26** | 0.33** | | |
| L IPS | -0.07 | -0.04 | 0.01 | 0.41** | 0.27** | 0.08 | 0.03 | 0.10 | -0.07 | 0.36** | 0.02 | 0.02 | |
| L Broca's | -0.08 | -0.08 | -0.06 | 0.28** | 0.25* | 0.12 | 0.01 | 0.03 | -0.06 | 0.18* | -0.04 | 0.00 | 0.39** |

**Fixation task with letters**

| | L V1 | L V4 | L LO | L VWFA-1 | L VWFA-2 | L text-mfs | L FFA-1 | L FFA-2 | R V1 | R VWFA-1 | R FFA-1 | R FFA-2 | L IPS |
|---|---|---|---|---|---|---|---|---|---|---|---|---|---|
| L V1 | | | | | | | | | | | | | |
| L V4 | 0.22** | | | | | | | | | | | | |
| L LO | 0.14 | 0.29** | | | | | | | | | | | |
| L VWFA-1 | 0.02 | 0.10 | 0.14* | | | | | | | | | | |
| L VWFA-2 | 0.04 | 0.07 | 0.01 | 0.45** | | | | | | | | | |
| L text-mfs | 0.01 | 0.09 | 0.04 | 0.40** | 0.28 | | | | | | | | |
| L FFA-1 | 0.10 | 0.24* | 0.27** | 0.30** | 0.20* | 0.28* | | | | | | | |
| L FFA-2 | 0.04 | 0.12 | 0.14 | 0.30** | 0.31** | 0.31 | 0.32** | | | | | | |
| R V1 | 0.28* | 0.13 | 0.16 | 0.03 | -0.02 | 0.11* | 0.11 | 0.10 | | | | | |
| R VWFA-1 | 0.07 | 0.12 | 0.16* | 0.44** | 0.23* | 0.14 | 0.15 | 0.14 | 0.08 | | | | |
| R FFA-1 | 0.07 | 0.21 | 0.20* | 0.18** | 0.14 | 0.16 | 0.37** | 0.28** | 0.07 | 0.33** | | | |
| R FFA-2 | 0.05 | 0.11 | 0.09 | 0.17* | 0.21 | 0.23* | 0.20* | 0.29* | 0.02 | 0.14 | 0.33** | | |
| L IPS | -0.00 | 0.12 | 0.02 | 0.31** | 0.26* | 0.19* | 0.07 | 0.13* | -0.05 | 0.20* | 0.02 | 0.09 | |
| L Broca's | 0.00 | 0.05 | -0.06 | 0.26** | 0.23* | 0.22** | 0.05 | 0.11 | -0.06 | 0.13 | -0.00 | 0.10* | 0.42** |

**Fixation task with shapes**

| | L V1 | L V4 | L LO | L VWFA-1 | L VWFA-2 | L text-mfs | L FFA-1 | L FFA-2 | R V1 | R VWFA-1 | R FFA-1 | R FFA-2 | L IPS |
|---|---|---|---|---|---|---|---|---|---|---|---|---|---|
| L V1 | | | | | | | | | | | | | |
| L V4 | 0.17* | | | | | | | | | | | | |
| L LO | 0.04 | 0.27** | | | | | | | | | | | |
| L VWFA-1 | -0.05 | 0.09 | 0.19** | | | | | | | | | | |
| L VWFA-2 | -0.01 | 0.03 | -0.03 | 0.40* | | | | | | | | | |
| L text-mfs | -0.05 | -0.01 | 0.13 | 0.35** | 0.24* | | | | | | | | |
| L FFA-1 | 0.04 | 0.24** | 0.21** | 0.28** | 0.20** | 0.28** | | | | | | | |
| L FFA-2 | -0.07 | 0.08 | 0.10 | 0.28** | 0.32** | 0.26* | 0.31** | | | | | | |
| R V1 | 0.25 | 0.25 | 0.25 | 0.02 | 0.05 | 0.01 | 0.09 | -0.01 | | | | | |
| R VWFA-1 | -0.01 | 0.07 | 0.19** | 0.39** | 0.28* | 0.22** | 0.12 | 0.16 | 0.04 | | | | |
| R FFA-1 | -0.05 | 0.18* | 0.20* | 0.19* | 0.16 | 0.17 | 0.39** | 0.24** | 0.10 | 0.32* | | | |
| R FFA-2 | -0.09 | 0.01 | 0.03 | 0.07 | 0.15 | 0.13 | 0.15 | 0.22 | -0.08 | 0.19* | 0.35** | | |
| L IPS | -0.02 | -0.00 | -0.01 | 0.38** | 0.25* | 0.14 | 0.04 | 0.08 | -0.04 | 0.24** | -0.01 | 0.05 | |
| L Broca's | 0.01 | -0.06 | -0.12 | 0.24** | 0.17 | 0.09 | -0.04 | -0.01 | -0.05 | 0.08 | -0.09 | 0.04 | 0.39** |

r: 0.5, 0.4, 0.3, 0.2, 0.1, 0, -0.1, -0.2, -0.3, -0.4, -0.5

**Figure S4: functional connectivity, related to Figure 5.** (**A**) Mean functional connectivity with left VWFA-1 as the "seed" region (In Fig. 5, Broca's was the seed). Before averaging, each subject's data were smoothed with a 2D Gaussian kernel (full-width at half-maximum = 5 mm). The data are masked to show only vertices where the correlation in trial-to-trial fluctuations was significant ($p$<0.05, corrected for false discovery rate), peaking at r>=0.4. (**B**) Correlations between trial-to-trial response variability in pairs of 14 regions, for trials when letters were presented in each task and stimulus condition. '**' indicates $P$<0.01 for a t-test comparing the mean correlation coefficient to 0. '*' indicates p<0.05. $P$-values corrected for false discovery rate.

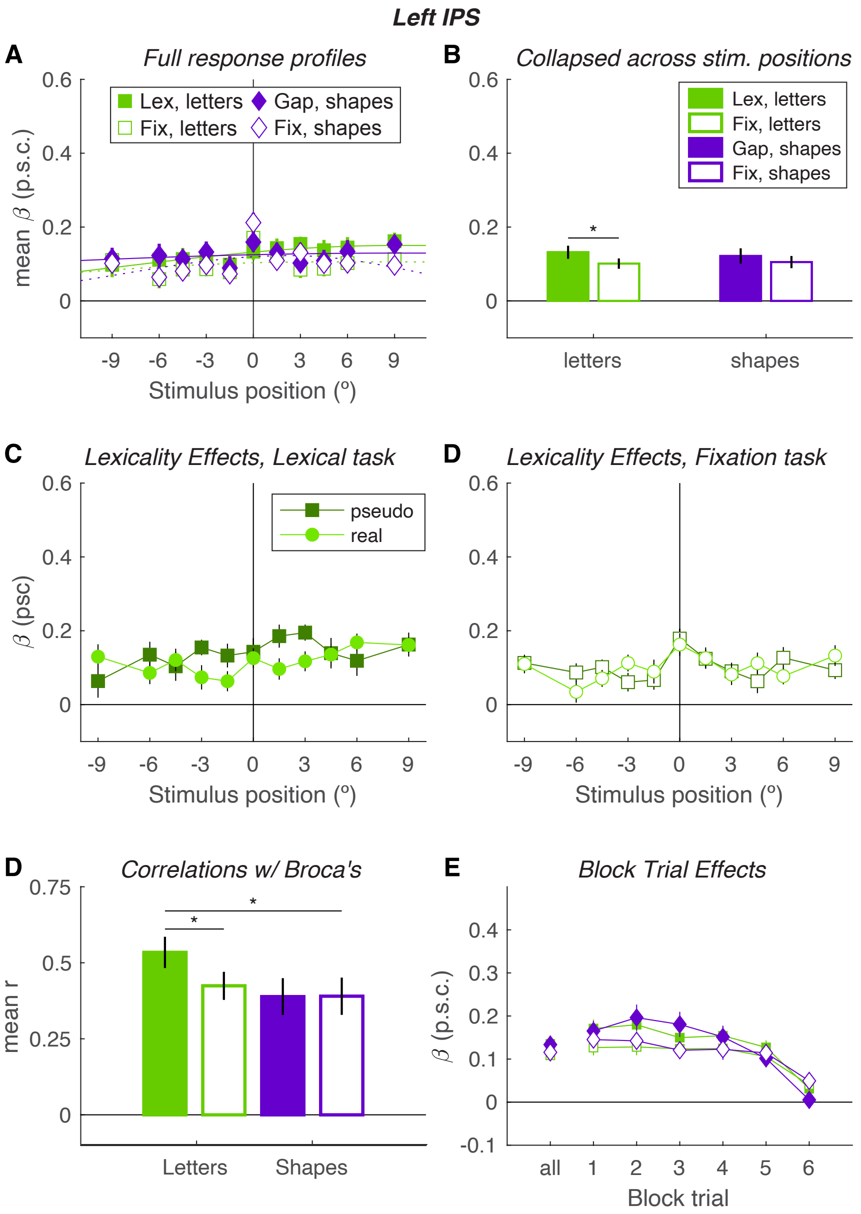


**Figure S5: activity in left IPS, related to Figures 2-5.** This left IPS region was defined in fsaverage space where the correlation with Broca's area r>0.2 (see Figure 5A). Each panel corresponds to a plot of VWFA activity in the main text. **A:** Although these spatial profiles appear relatively flat, there was a significant main effect of absolute eccentricity (*P*=0.02), which was larger for letters than shapes (*P*=0.03). **B:** There was a main effect of task: "attend-stimuli" > "attend-fixation" (*P*=0.003), which did not interact with stimulus type (*P*=0.31, BF=0.42). **C:** In the lexical decision task, but not in the fixation task, there was a significant effect of lexicality (pseudowords>real words; *P*<0.001) that interacted with eccentricity (*P*=0.02) **D:** The functional connectivity with Broca's area was stronger when letters were presented than shapes (*P*=0.003). That effect interacted with task (*P*=0.04, BF=1.6), with a task effect (lexical > fixation) only for letters (*P*=0.01, BF=4.2). **E**: As in other areas, the overall BOLD response decreased across trials in each block ($P=3\times10^{-29}$), and the task effects diminished as well (*P*=0.002). We also analyzed data in a more posterior region that encompassed IPS-0, IPS-1 and IPS-2, defined from the Wang et al (2015) atlas (not shown). Its activity was similar to what is pictured here, but with a stronger task effect (gap>fixation) in the BOLD response to shapes.

| REAL WORDS | | | | | PSEUDOWORDS | | | |
|---|---|---|---|---|---|---|---|---|
| able | form | line | says | | aden | frur | muna | thir |
| also | four | live | seem | | aizi | futh | muro | thok |
| away | free | long | seen | | alst | gomy | neee | thom |
| back | full | look | show | | alur | grur | neng | tian |
| best | gave | lord | side | | aror | haca | neny | tirl |
| body | girl | lost | some | | awan | hece | nery | tith |
| book | give | love | soon | | awat | huso | okis | toto |
| both | gone | made | sort | | blso | inen | onch | trur |
| call | good | make | such | | blus | iney | onem | unth |
| came | hair | many | sure | | cery | inld | oner | upod |
| case | half | mean | take | | cich | inlk | onld | upom |
| cent | hand | mind | talk | | cror | inll | onll | usen |
| city | hard | miss | tell | | dasy | inly | onlp | vech |
| come | head | most | than | | doch | inod | onod | veey |
| dark | held | much | then | | eady | inom | onom | vefe |
| days | help | must | time | | edey | inow | onot | vith |
| does | here | name | told | | egen | inth | onow | wepy |
| done | high | need | true | | eger | itis | onte | wete |
| door | home | next | took | | elsk | juch | onth | wher |
| down | idea | once | turn | | elst | juld | onty | wier |
| each | just | only | upon | | elur | kner | oved | wiey |
| else | keep | open | used | | enth | knly | ovek | wifa |
| even | kept | over | very | | evew | knto | ovem | wiky |
| ever | kind | part | view | | evey | lelk | ovep | wiso |
| eyes | knew | past | want | | eype | lely | ovew | wito |
| face | know | play | week | | faly | lery | rele | woto |
| fact | land | poor | well | | fito | lith | roso | yeen |
| feel | last | read | went | | frar | lito | saly | yeey |
| feet | late | real | wife | | frem | mely | siey | yery |
| felt | left | rest | word | | frer | mito | sirm | yexi |
| find | less | road | work | | fris | moey | soto | yext |
| five | life | room | year | | frme | morr | tagh | yont |
| food | like | same | your | | fror | mugh | thia | yous |

**Table S1: Lists of real words and pseudo used in the main experiment. Related to Figure 1A.**
